# Development of a Two-Stage Life Cycle Model to Inform the Trap and Haul Program for Coho Salmon in the Lewis River, Washington

**DOI:** 10.1101/2025.04.30.651546

**Authors:** John M. Plumb, Russell W. Perry

## Abstract

Restoration of salmon populations in the upper Lewis River Basin depends on a trap-and-haul program owing to the Lewis River Hydroelectric Project (Project) operated by PacifiCorp and Cowlitz PUD (Utilities), which has been a barrier to salmon passage since the 1930s. Thus, sustaining the Coho salmon (*Oncorhynchus kisutch*) population upstream of the Project currently depends on two fundamental factors: (1) the collection of upstream migrating adult Coho salmon at Merwin Dam, the lower most dam within the Project, and transporting them by truck to spawn above Swift Dam, the upper most dam within the Project; and (2) the collection of out-migrating juvenile Coho salmon at the downstream collection facility at Swift Dam for transport and release below the Project. The reintroduction program began once the downstream collection facility at Swift Dam was commissioned in late-2012 with the first year of transport data being collected in 2013. Over the past decade, the Utilities have been collecting data on juvenile outmigrants and adult fish returns at the dams. The need to construct a life cycle model for Lewis River anadromous fish was identified by the Lewis River Aquatic Technical Subgroup, with the understanding that many years (>15) of data collection are needed to adequately measure the life cycle production of coho salmon. Use of past data to construct models could help inform future data collection and provide a framework that can be updated annually to measure trap and haul program performance within a life cycle context (Note: Data are not currently available from PacifiCorp. Contact organization <u>Chris Karchesky</u> for further information).

Because Coho salmon can live as long as five years, estimating demographic parameters for Coho salmon populations over their life cycle requires at least 10 or more years of data collection. Over the past decade, PacifiCorp has been collecting data on fish collection efficiency and the numbers of adult and juvenile salmon transported around the Lewis River dams, providing sufficient data to formulate a life cycle model that can guide future data collection efforts and provide preliminary information to resource managers The goal of the statistical life cycle model was to estimate annual production and survival during two critical life-stage transitions (1) the freshwater production from escapement of adults released upstream of Swift Dam, and the collection of downstream migrating juveniles at the passage facility at Swift Dam, and (2) the smolt-to-adult survival from the time of collection at Swift Dam to their return as adults. We used the Beverton-Holt stock-recruitment model to estimate juvenile production from the number of spawners. This approach allowed us to test for density dependence at current spawner abundances while estimating annual productivity, defined as the number of juveniles produced per spawner at low spawner abundance. Productivity was then expressed as a function of the number of juveniles collected and transported downstream of the Project. Because juvenile Fish Collection Efficiency (FCE) directly affects the number of juveniles that survive to continue downstream migration, FCE is a primary determinant of fish production. Consequently, the modeling framework is well suited to evaluate the performance of trap and haul programs within a life cycle context.

The objectives of this study were to: (1) gather and collate available data on adult and juvenile Coho salmon at Merwin and Swift dams, (2) quantify adult escapement, juvenile abundance, and the age at outmigration and adult return, (3) describe, formulate, and fit the integrated population model (IPM) to the data, and (4) summarize our findings, identify data gaps, and identify potential opportunities for future studies that could provide information used to improve model estimation and inference. Our key findings were: (1) over and above the number of spawning females, FCE was the primary factor affecting productivity of Coho salmon above Swift Dam, (2) smolt-to-adult return (SAR) rates were relatively high considering that harvest was included in the estimate, averaging about 4.5% and ranging as high as 12.9%, and (3) juvenile capacity upriver of Swift Dam was difficult to estimate due to the limited range in spawning females over the time series of data, suggesting the model may be improved by collecting data at higher spawner abundances. In addition, by including FCE in the model, we estimated that the median pre-collection productivity, defined as the number of juveniles produced per spawner when FCE = 1, was 64 juveniles per spawner. Because this two-stage life cycle model partitions factors that affect fish production in river versus the ocean, the model estimates should help inform fishery managers about the overall role that fish collection at Swift Dam plays in the recovery and sustainability of Lewis River Coho salmon. By providing the model with (1) more years of data, (2) higher numbers of spawning females, and (3) data on age at juvenile migration in relation to age at adult return greater certainty in the estimates of capacity and SAR can be attained. Ultimately, information provided by the model can assist in the evaluation and continued improvement of the current trap and haul program to support anadromous fishes in the Lewis River Basin.

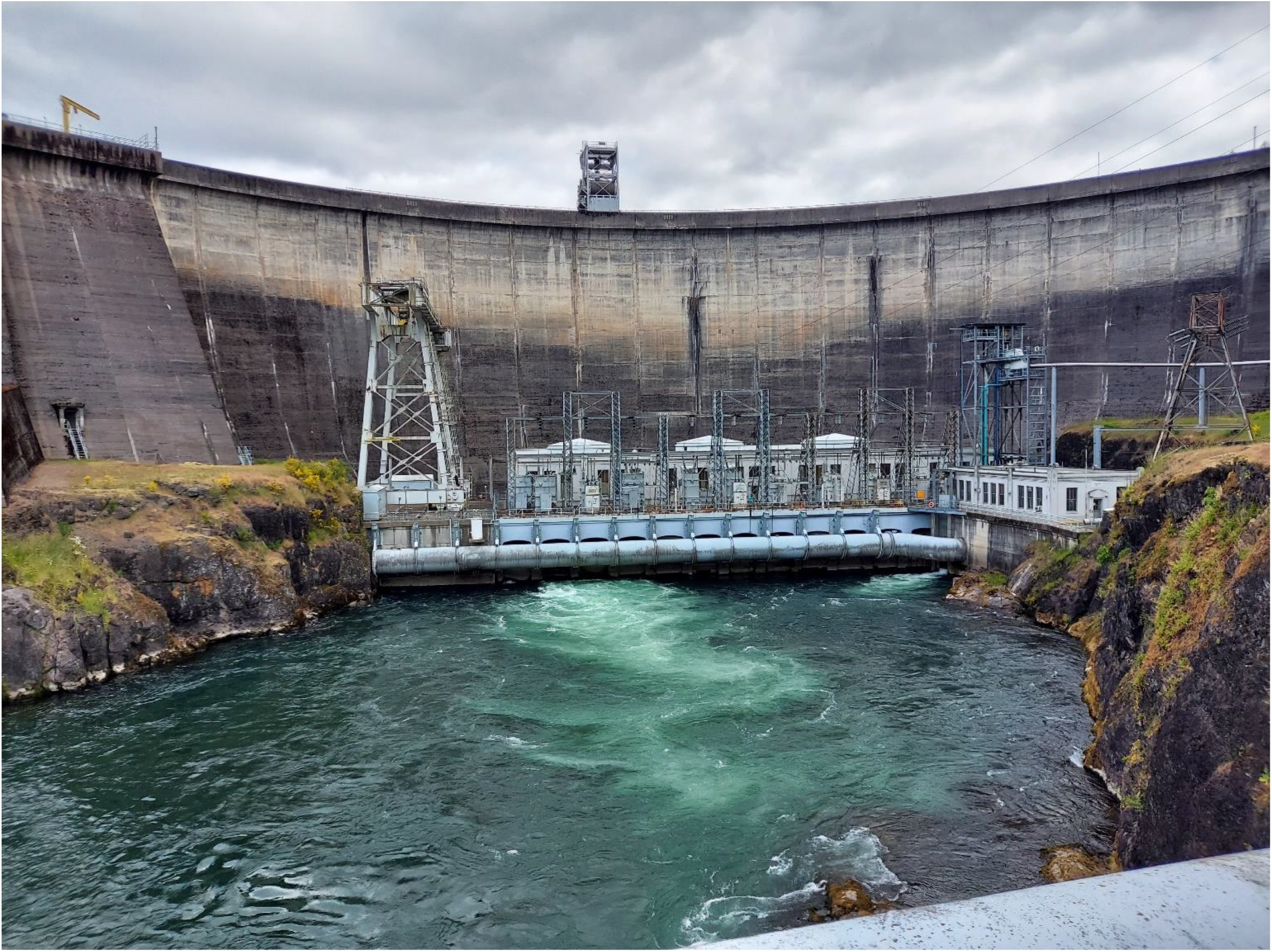

**Cover.** Looking upstream on the Lewis River at Merwin Dam, Washington, August, 2024. Photograph by John Plumb, U.S. Geological Survey.

**Conversion Factors:** 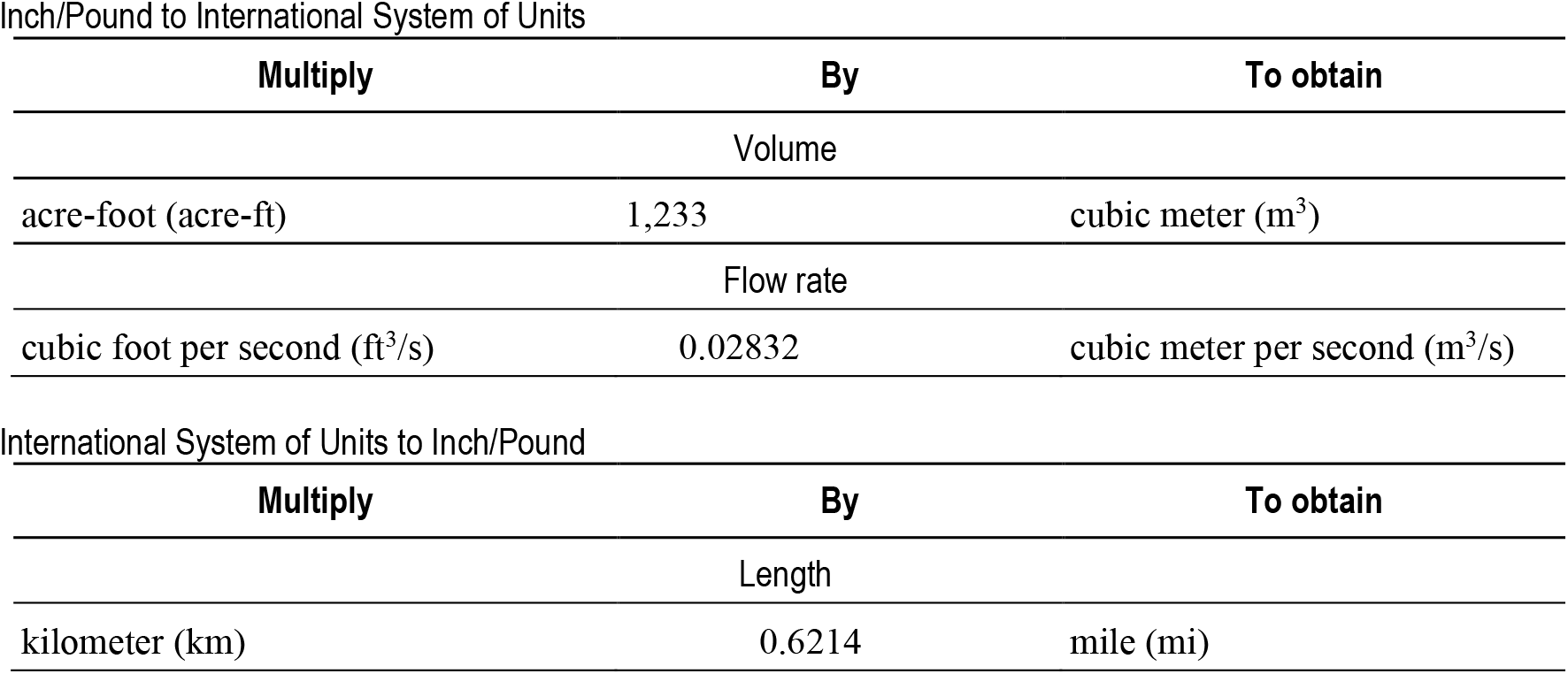

## Introduction

Herein, we present a two-stage state-space life cycle model (or Integrated Population Model; IPM) for Coho salmon (*Oncorhynchus kisutch*) in the Lewis River (fig 1.). The IPM is intended to complement existing spreadsheet-based deterministic life cycle models often used by fisheries managers to track demographic parameters and determine appropriate hatchery production. In contrast to the deterministic spreadsheet model, the IPM approach incorporates both observation and process error in abundance and production estimates (Gelman and others, 2004), allowing users to estimate both mean and annual life-cycle demographic parameters and associated uncertainty. Our approach is an extension of the single-stage state-space framework first developed by Fleischman and others (2013) and applied by Courter and others (2019) to the Clackamas River steelhead (*O. mykiss*) population, representing a variation of the two-stage state-space model that we applied to Coho salmon in the Cowlitz River Basin, Washington (Plumb and Perry, 2020). The state-space model consists of two parts: (1) a process model for the underlying state dynamics, and (2) an observation model that links the data to the true underlying state. The state-space model may also be thought of as a hierarchical model where the state (abundance of fish) evolves according to a dynamic population process model (e.g., Beverton-Holt or Ricker) with some process error, and observations on the state (“the data”) are made conditional on the true but unobservable state (Beverton and Holt Ricker 1954).

**Figure 1.**
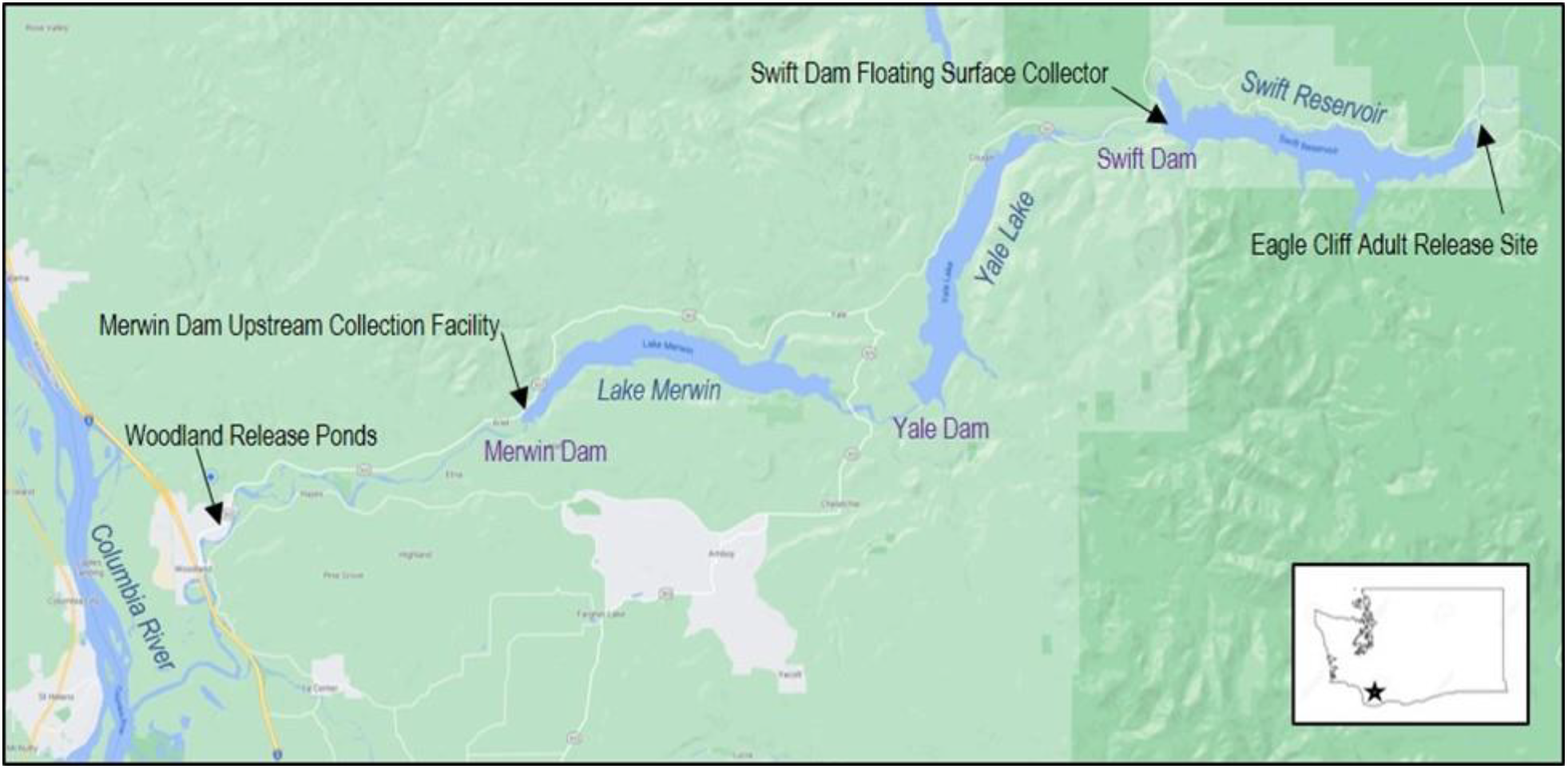
Map of the Lewis River Hydroelectric Project, showing dams, reservoirs, and locations important to the trap and haul program for salmonids.

The modeling approach first breaks the life cycle into juvenile and adult life stages and then includes population composition, which is comprised of sex, rearing origin (hatchery or naturally produced), and juvenile and adult age structure. Multistage life cycle models provide a powerful analytical framework for understanding how each life stage of a population contributes to population growth rate (Moussalli and Hilborn 1986; Greene and Beechie 2004). Multistage models may also be used as an analytical framework to explicitly estimate demographic parameters of a population model. Unlike single-stage stock-recruitment models, multistage models allow population growth rates to be partitioned among life stages rather than aggregated over an entire life cycle. Partitioning among life stages facilitates estimation of (1) stage-specific density dependence, and (2) stage-specific effects of environmental factors or management actions. For example, Zabel and others (2006) estimated parameters of a multistage model used in the context of a population viability analysis for spring/summer Chinook salmon (*Oncorhynchus tshawytscha*) in the Snake River.

Typically, data informing estimates of abundance at particular “check points” in the life cycle determines the complexity of the multistage model that can be fit to the data. For Coho salmon in the Lewis River Basin, we have developed a two-stage model that encompasses: (1) upstream passage of spawners at Merwin Dam to the subsequent downstream passage of their progeny at Swift Dam, and (2) downstream collection of juveniles at Swift Dam to their subsequent return from the ocean and passage at Merwin Dam 1−3 years later. This approach partitions the life cycle of Coho salmon both spatially and temporally, which allows us to fit and compare alternative models with covariates specific to each stage. This report describes the structure of the two-stage life cycle model as applied to Lewis River Coho salmon, presents preliminary results from fitting the model to data, and outlines future directions and developments.

## Study Area

The Lewis River Hydroelectric Project (Project) begins approximately 16 km east of Woodland, Washington (fig. 1), and consists of four impoundments. The sequence of the four Lewis River impoundments upstream of the confluence of the Lewis and Columbia Rivers is: Lake Merwin, Yale Lake, Swift Reservoir, and Swift Number Two Forebay. Currently, reintroduction efforts for anadromous salmonids are occurring in habitats upstream of Swift Dam. The target species identified for reintroduction are spring Chinook salmon (*O. tshawytscha*), Coho salmon, and winter Steelhead (*O. mykiss*). As part of this program, adult salmonids are collected at Merwin Dam and then transported by truck and released into Swift Reservoir to continue their migration to the upper watershed of the Lewis River. When juvenile salmonids migrate downstream, they are collected at the collection facility at Swift Dam, referred to as the Swift Floating Surface Collector (Swift FSC) located at the downstream extent of Swift Reservoir. From there, the juveniles are transported by truck to below Merwin Dam and released at the Woodland Release Ponds before volitionally reentering the Lewis River to continue their downstream migration (fig. 1). Because there is no fish passage currently at Yale or Merwin Dams, juvenile fish that pass Swift Dam into Yale Lake are currently considered lost to the anadromous population.

## Methods

Life cycle models can range from very simple theoretically based population models (e.g., the Beverton-Holt or Ricker stock-recruitment models) to very complex spatially explicit simulation models linked to hydrosystem hydrodynamic models (e.g., the COMPASS model for a single transition in a life cycle model, Zabel and others, 2008). We chose to develop a model of intermediate complexity that casts the two-stage life cycle model in a state-space framework (Newman and others, 2014). We chose to use a state-space framework implemented in a Bayesian context because:

- The model provides both a statistical estimation framework for retrospective statistical analysis and a stochastic simulation framework for prospective analysis to evaluate alternative management actions.
- The model accounts for uncertainty in abundance estimates. A state-space framework accounts for observation uncertainty in the abundance estimates and other data (e.g., age structure) while simultaneously estimating process uncertainty.
- The model allows for missing data. By drawing missing data from an appropriate probability model, uncertainty owing to missing data can be propagated without having to omit data or assume fixed values for missing data.

Thus, a two-stage state-space life cycle model for Coho salmon provides an appropriate balance between model complexity, tractability, and applicability given the goals of performing both retrospective and prospective analysis to guide future management.

### Life Cycle Structure of Lewis River Basin Coho salmon

The state-space model consists of two parts: (1) a process model for the underlying state dynamics, and (2) an observation model that links the data to the true underlying state. The state-space model may also be thought of as a hierarchical model where the state (abundance) evolves according to a process model for population dynamics (e.g., a Beverton-Holt model) with some process error, and observations on the state (“the data”) are made conditional on the true but unobservable state.

First, to model the adult to juvenile transition (fig. 2), we used the Beverton-Holt model to express the number of juveniles collected at Swift Dam as a density-dependent function of the number of female adults passing Swift Dam in brood year *y* (Beverton and Holt 1957):

**Figure 2.**
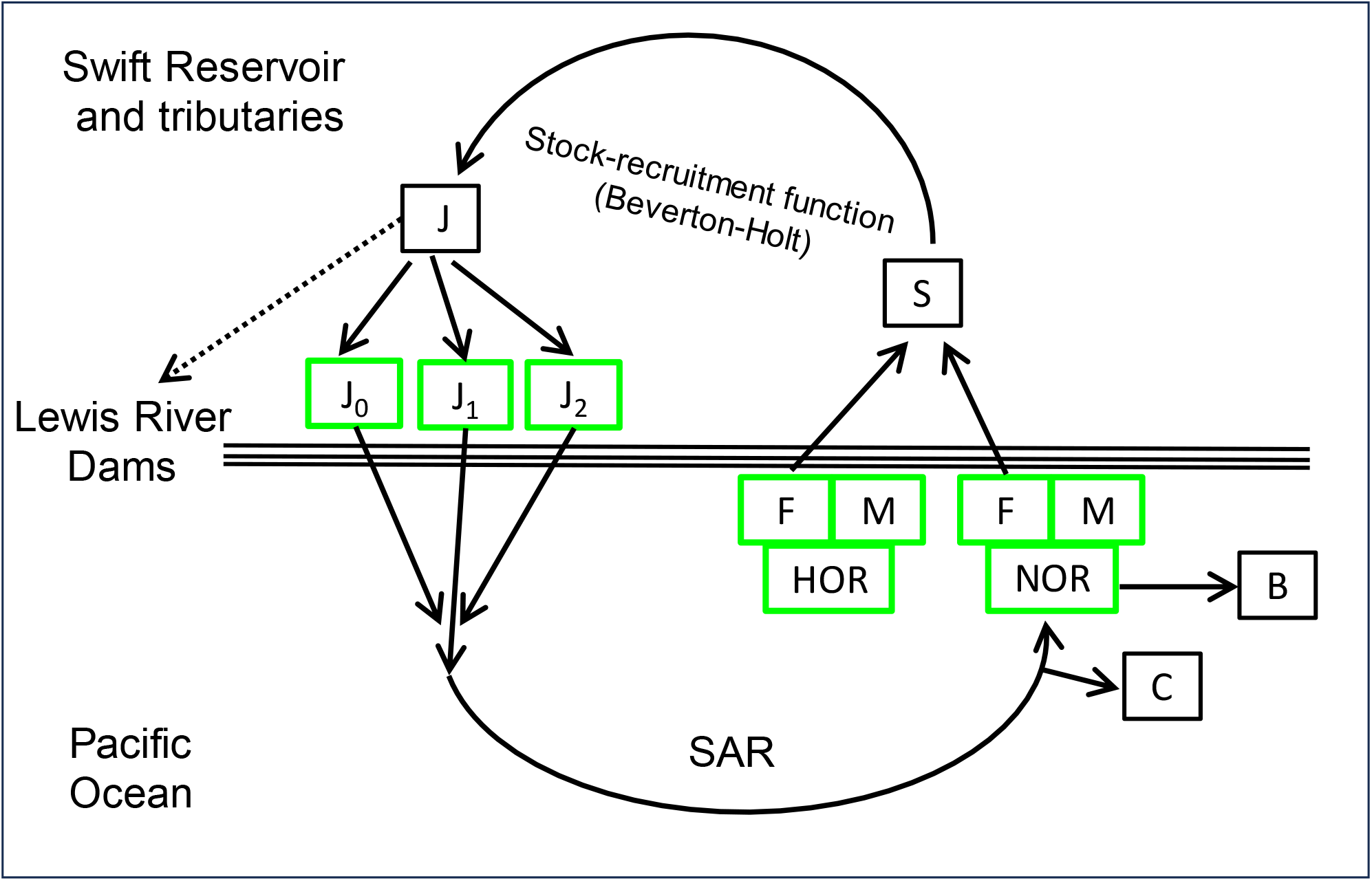
Schematic illustrating the current structure of the two-stage life cycle model for Lewis River Coho salmon, showing juvenile age-structure at Swift Dam (J_0_ = subyearling, J_1_ = yearling, J_2_ = Two year-old outmigrants), and sex (F = female, and M = male) and origin (HOR = hatchery, NOR = natural) of spawning adults, while accounting for harvest (C) and brood stock removal (B). The spawner-to-juvenile transition is modeled using a stock-recruitment function, and the juvenile-to-adult transition is modeled with a smolt-to-adult return rate (SAR). Solid arrows designate the structure that is currently incorporated into the model. The dashed arrows designate unknown quantity of fish that passed downstream of Swift Dam and lost to the anadromous population. Note that age structure of returning females and males is only partially known from the jack return of males and therefore is not shown.

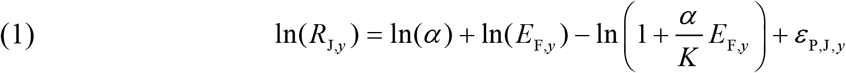

Where:

*R*_J,*y*_ is the true but unobserved number of collected juvenile recruits produced by

*E*_F,*y*_ female adult escapement in brood year *y*;

*α* is the productivity parameter estimating the slope at the origin (juvenile recruits per female spawner at low spawner abundance in the absence of density dependence); and K is carrying capacity of juvenile recruits and is defined as the maximum number of juveniles that can be produced upriver of Swift Dam. The process error term, *ε* _P,J, *y*_, quantifies annual variation in productivity driven by environmental variation with mean of zero and standard deviation *σ* _P,J_ . Because productivity of juveniles directly depends on the fraction of juvenile Coho salmon collected at Swift Dam, we used annual estimates of fish collection efficiency (FCE), which measures fish capture relative to the number of fish that approached the floating fish collector. This provides an index of fish facility performance from year to year. We modeled the productivity parameter *α* as function of the annual FCE and other covariates for Coho salmon in the following manner:

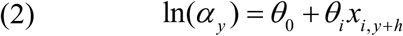

Where:

exp (*θ*_0_)is the mean productivity at the mean covariate value, *θ*_*i*_ is the slope for the effect of the *i*th covariate on annual productivity, and

*x*_*i*, *y* + *h*_ is the *i*th covariate with an offset of *h* years on brood year to reflect the life stage in which the covariate is hypothesized to most affect productivity.

The proportion of early spawning Coho salmon was defined as those adult Coho salmon that arrive at Merwin Dam and are transported during the early (September) compared to the late (October) portion of the spawning migration. Note because the stock-recruitment model was informed by just 10 years of data, we did not fit the model using all covariates on productivity simultaneously. Rather we used a stepwise approach where *x*_2_ − *x*_5_ were respectively used to estimate *θ* _2_ − *θ*_5_ within separate models that also included the effect of FCE with *θ*_1_ *x*_1, *y* + 2_ . Specifically, *x*_2, *y* +0_ was the data vector for the proportion of hatchery-origin female spawners, *x*_3, *y* + 0_ was the proportion of early spawners (*Ep*), and *x*_4, *y* +0_ was the maximum monthly winter flow indexed to brood year *y*. The minimum summer flows, *x*_5, *y* + 2_, were indexed to the yearling outmigration in year *y*+2. Flow data were obtained from the U.S. Geological Survey (USGS) streamgage (USGS station number14216500; U.S. Geological Survey, 2024) on the Muddy River located at the upper end of Swift Reservoir. Importantly, when drawing inferences about the effect of FCE on productivity or SAR, we used the model that only included the effect of FCE on productivity in the model. Although the model’s framework is designed to accommodate covariates associated oceanic effects on SAR, we refrained from adding covariates on SAR at this time due to the lack of information linking juvenile age at outmigration to adult age at return to Merwin Dam.

For the juvenile to adult transition, we model the number of adult returns as a lognormal function of a density independent smolt-to-adult return rate (SAR):

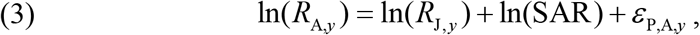

where

*R*_A, *y*_ is the number of adult recruits (male and female) produced from the *R*_*J*,*y*_ juveniles collected at Swift Dam that arose from female spawners in brood year *y*; and

*ε* _P,A,*y*_ is a normally distributed process error with mean zero and standard deviation *σ* _P,A_ .

Given that juveniles can pass Swift Dam at ages 0-2 and adults return at ages 1-3, the initial two years of juvenile recruits and 3 years of adult recruits that produced returns beginning in 2013 were not linked to the two-stage stock-recruitment model. These initial state vectors were estimated as draws from common log-normal distributions with parameters ln (*R*_J,0_), *σ* _RJ0_ and ln (*R*_A,0_), *σ* _RA0_, respectively for juveniles and adults.

Given juvenile recruits, the number of juveniles *a*_J_ passing Swift Dam in calendar year *t* was modeled as:

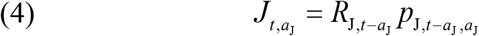

where

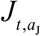 is the number of juveniles in year *t* of age *a*_J_ ; and 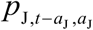 is the proportion of juvenile recruits from brood year *y* = *t* − *a*_J_ emigrating at age *a*_J_ (fig. 3).

**Figure 3.**
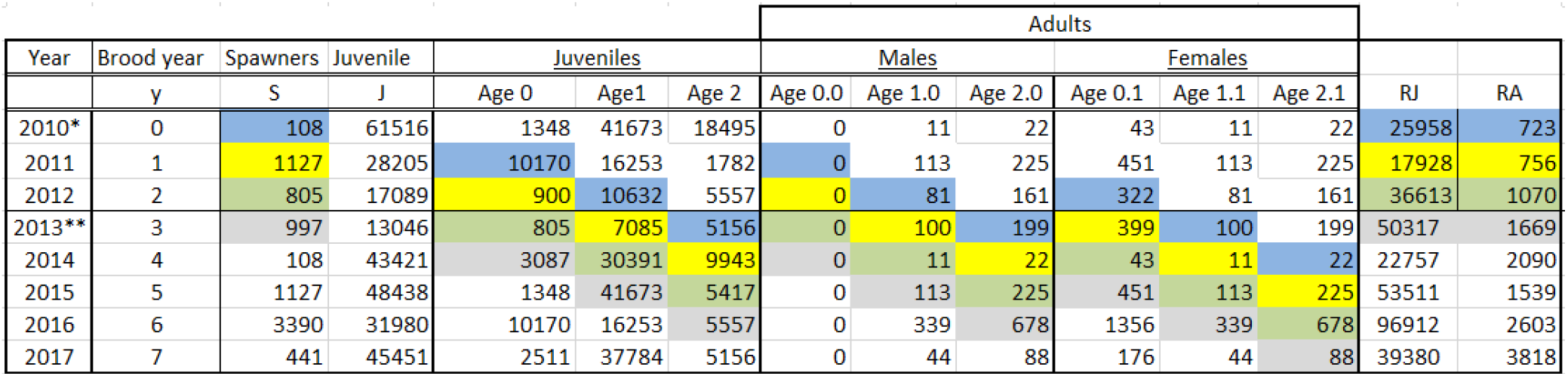
Illustration using hypothetical data to reconstruct age-specific juvenile abundance and ocean age-specific adult returns conditional on juvenile age at outmigration from recruits generated by spawners in brood year *y*.

Given adult recruits, we model the number of adults returning at age *a*_A_ outmigration strategy *o* as:

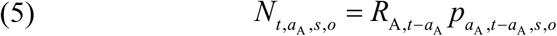

where

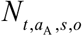 is the number of adults returning in year *t* at age *a*_A_, of sex *s* and outmigration strategy *o* and 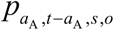 is the proportion of the recruits from brood year *y* = *t* − *a*_A_ returning at *a*_A_, sex *s*, and outmigration strategy *o* (fig. 3).

Outmigration strategy refers to whether juveniles were collected at Swift Dam as subyearlings, yearlings, or two-year-old outmigrants. Because data on the age structure of returning adults for a given age at juvenile outmigration are absent, our current formulation of the two-state model does not account for juvenile age at outmigration in the adult returns, but in the future such information would allow us to estimate SAR separately for each outmigration strategy. Nonetheless, given three adult ocean age classes (MacLellan and Gillespie, 2015; fig. 3), and three outmigration strategies, the brood-year specific return probabilities are 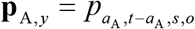.

We model the brood-year composition using this structure for several reasons. First, juvenile recruitment is expressed as a function of female escapement, requiring estimation of sex structure. Second, age 0 – 2 juvenile proportions are of direct interest. Third, both jack and adult abundances have been recorded since 2012, whereas comprehensive age structure data has not yet been collected. Thus, utilizing both the jack and adult abundance data requires estimating the proportion of jacks in the brood year return.

Vectors of brood-year-specific marginal and conditional probabilities were modeled hierarchically as draws from a Dirichlet distribution (Fleischman et al. 2013, Scheuerell et al. (2021):

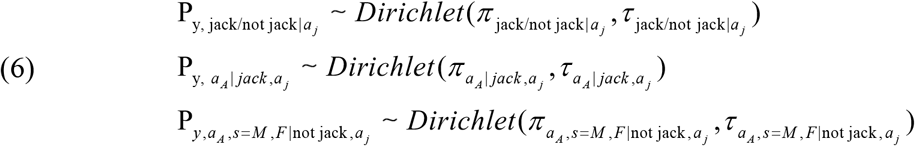

where *π* is the vector of mean proportions in each age, sex, and age at ocean entry category, and is an inverse variance parameter. This model structure allows for different levels of interannual variability for each set of probabilities. This hierarchical structure can be used for accommodating missing age-structure data. That is, for years with missing age structure, the model imputes **π** τ.

Given age-specific juvenile emigration and adult returns from brood year *y* in calendar *t*, the total number of juveniles passing Swift Dam in calendar year *t* is the sum of the abundance-at-age over all ages:

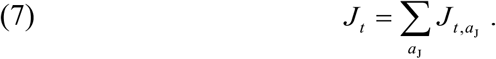

Note that the current model for Coho salmon at Swift Dam accounts for brood stock removal but does not account for juvenile age at outmigration or harvest rates; however, we show here how these estimates can be incorporated into the current framework of the model for future applications. For example, the total number of returning adults in year *t* is the sum of the abundances across all age, sex, and outmigration classes:

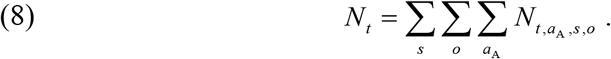

Returns of subclasses in year *t* are calculated similarly by summing over the appropriate indices. For example, the number of female returns in year *t* is

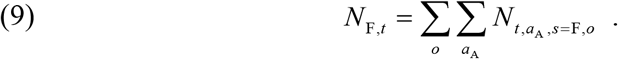

In addition, the age, sex, and outmigration structure in the adult returns each calendar year is

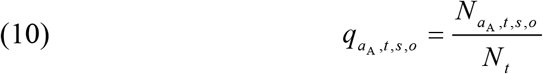

where 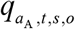 is the fraction of the adults returning in year *t* at age *a*_A_, sex *s*, and outmigration strategy *o* (i.e., if data on outmigration strategy were included in the model).

Escapement of spawners past Swift Dam in calendar year *t* includes hatchery-origin adults released upriver of Swift Dam to spawn in the wild (*H*_*t*_) plus naturally produced spawners (*N*_*t*_). The model can accommodate those naturally produced spawners that survive harvest (*C*_*t*_) or those taken for hatchery brood stock (*B*_*t*_):

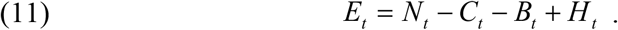

The annual harvest rate below Swift Dam can be a composite of ocean harvest that is assumed constant across ages and sexes of returns each year such that catch is:

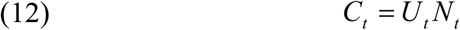

where *U*_*t*_ is the composite harvest rate representing the fraction of returns that were harvested in year *t*.

Escapement of female spawners, which dictates juvenile recruits, is:

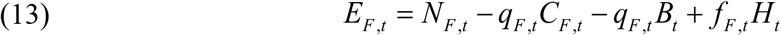

where *f* _*F*,*t*_ is the fraction of hatchery-origin adults that are female and *q*_*F*,*t*_ is the proportion of females in naturally produced returns, calculated as:

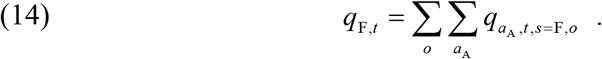

### The Observation Models

Observations to inform the parameters of the state model include:

1. Catch of age 0, 1, and 2 outmigrants collected annually at the Swift FSC during 2013– 2023;
2. Census counts of annual abundance of hatchery-origin and naturally produced adults released above Swift Dam for 2013–2023;
3. Estimates of annual run composition of hatchery and naturally produced adults in terms of jacks, adults, and sex, 2013–2023;
4. Estimates of the annual number of naturally produced adults removed for hatchery brood stock for 2013–2023; and
5. Estimates of ocean harvest rate indices as well as harvest above Swift Dam for 2013–2023 (not yet included in the model).

#### Juvenile Abundance

Estimates of abundance of naturally produced juvenile Coho salmon collected at Swift Dam were assumed to be lognormally distributed about the true abundance:

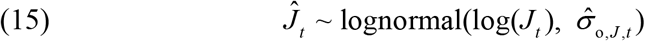

where *ĵ*_*t*_ is the total abundance estimate that includes age 0, 1, and 2 fish collected at Swift Dam in year *t*, and *J*_*t*_ is the true but latent state vector of juvenile abundances, 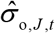 and is the observation error. In contrast to escapement, where we estimate observation error within the IPM, here we assume the observation error is known using annual estimates of uncertainty in the juvenile abundance
estimates calculated as 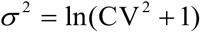, following Fleischman and others (2013).

The observation model for juvenile age composition was done by expressing the juvenile abundance estimates for age 0, age 1, and age 2 as a multinomial distribution:

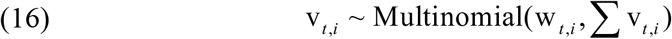

Where v_*t*,*i*_ is the vector of abundances for each age at outmigration entry in year *t*, w_*t*,*i*_ is a vector of probabilities for each category, and *i* indexes juvenile age at the time of collection at Swift Dam.

#### Adult Abundance

The census counts for adult escapement upstream of Swift Dam were assumed to be lognormally distributed with known error (CV=0.01) calculated as 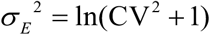, following Fleischman and others (2013):

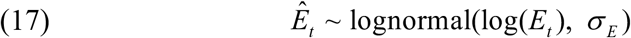

where *Ê* _*t*_ is the total escapement for jacks, males, and females released upriver of Swift Dam in year *t*, and *E*_*t*_ is the true but latent state vector of adult escapement, *σ* _*E*_ and is the observation error.

In the observation model for adult age composition, census counts of jacks, older males, and females were expressed as a multinomial distribution:

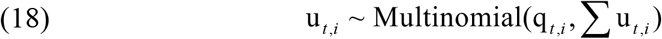

Where u_*t*,*i*_ is the vector of adult abundances collected at Merwin Dam by age-at-ocean entry (assuming this information was available), age at return, and sex category in year *t*, q_*t*,*i*_ is a vector of probabilities for each category, and *i* indexes jacks and older adults and sexes.

### Parameter Estimation

All parameters and unknown states were estimated in a Bayesian framework using JAGS software (Plummer 2009) as implemented through the ‘runjags’ package (Denwood, 2016) of the R statistical programming platform (R Core Team 2023). JAGS is a Bayesian estimation software package that implements Markov Chain Monte Carlo (MCMC) sampling using a Gibbs or Metropolis-Hastings sampler. Prior distributions for most parameters were set according to Fleischman and others (2013). However, preliminary model using a weakly informed prior distribution indicated the data were insufficient to estimate a capacity. Therefore, we used information from a meta-analysis on Coho salmon populations throughout the Pacific Northwest region to inform the prior distribution on capacity. We compare posterior distributions when using the weakly informed versus the informed prior distribution. We used the log normal distribution parameters provided by Korman and Tompkins (2014) (mean, μ = 7.46 and standard deviation, SD = 0.67) as the informed prior distribution on capacity. For the weakly informed prior distribution, we used a truncated normal distribution with a μ = 10,000 and SD = 1,000,000. Comparison of posterior distributions resulting from different prior distributions helped provide a range in capacity estimates for the estimated habitat upriver of Swift Dam. Importantly, the Korman and Tompkins (2014) estimates of capacity were provided on a per km basis. Therefore, we used 186.9 km of habitat upriver of Swift Reservoir and an additional 14.5 km for the reservoir resulting in a total of 201.4 km of habitat for Coho salmon. This assumes similar habitat carrying capacity between the reservoir and riverine habitats upriver of Swift Dam.

We ran three independent MCMC chains each for 80,000 iterations, discarding the first 30,000 to ensure each the chain had converged to its stationary stable distribution. We then thinned the final 50,000 iterations to 1 in 50 to reduce autocorrelation yielding a final sample of 1,000 draws from each chain. Convergence of each parameter was checked visually to ensure mixing of the chains, and quantitatively by ensuring that the Rubin-Gelman statistic 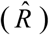 was less than 1.1 (Gelman and others, 2004).

### Estimation of Abundance of Juvenile Coho salmon Collected at Swift Dam

To implement the two-stage life cycle model, we first had to estimate the abundance of juveniles collected at Swift Dam. Of the total fish collected, a varying fraction of fish are sampled to determine size and age composition. Although most days 100% of the fish are sampled, on infrequent days during high fish passage, sample rates are lowered for logistical reasons. Over the time series, daily sample rates were as low as 0.1, but 87% of the time sample rates = 1. We used the daily (*d*) sample rates in year *t*, (*r*_*d*,*t*_) and the counts of juvenile salmon in the sample tank at Swift Dam (*c*_*d*,*t*_), to estimate the age-specific (*a*) total abundance of juveniles collected (*J*_*d*,*t*_) at the dam.

Estimating age-specific abundance requires classifying juveniles by age (0, 1, and 2). Catch-at-age was determined by PacifiCorp personnel, and we assumed no classification error in age assignment (fig. 4). Daily counts of age zero fish were considered a complete census, without classification error, owing to their small size at date. Thus, estimation of the total number of age 0 fish, *J*_0*d*,*t*_, involved simply inflating sample-tank counts by the sample rate, if counts, *c*_0*d*,*t*_, were binomially distributed:

**Figure 4.**
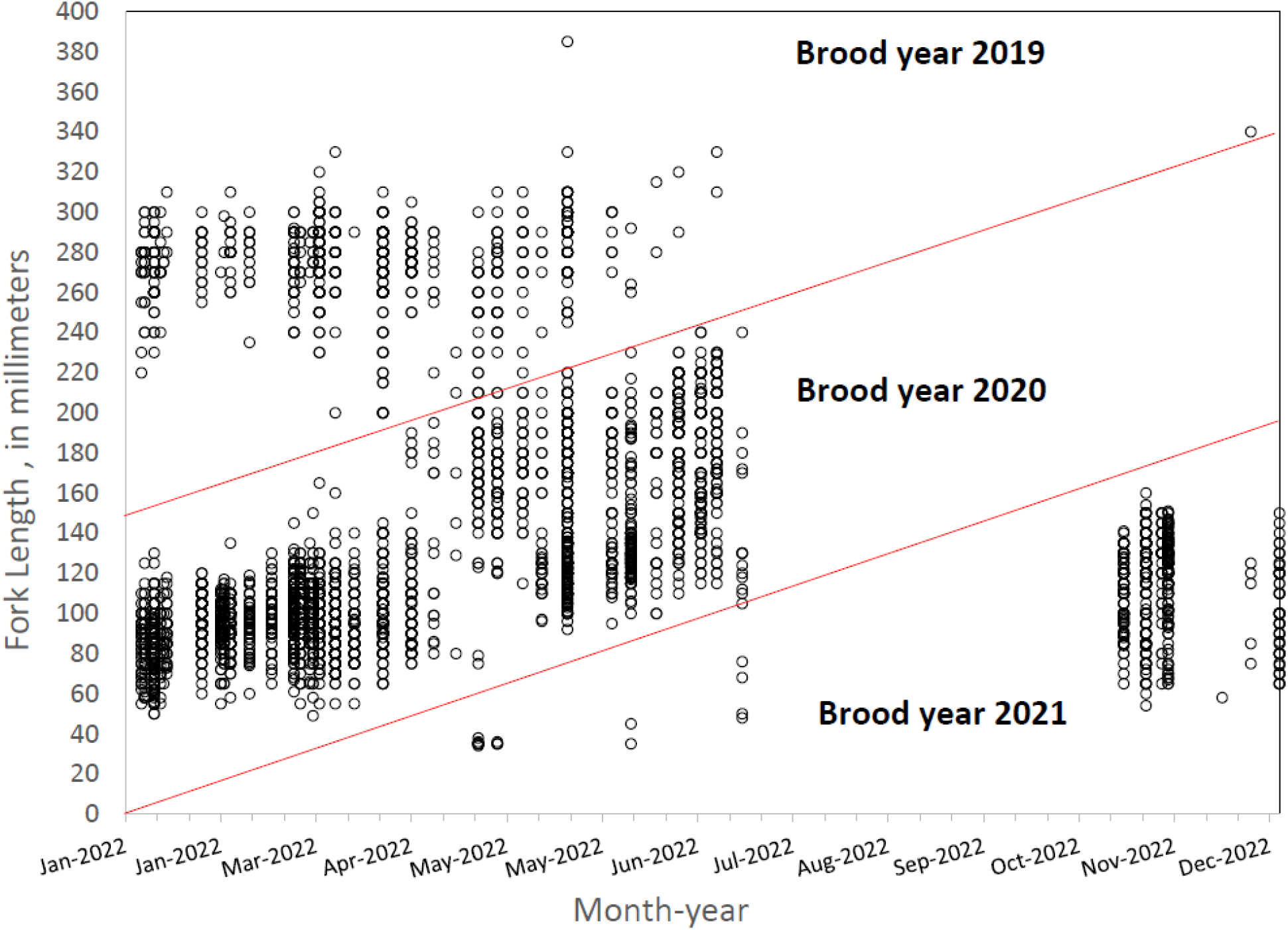
Example of age assignment to brood year (red lines) by fish size and month for juvenile Coho salmon collected at Swift Dam during 2022.

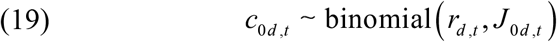

and for age 1,

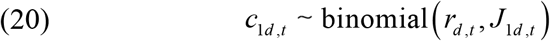

and age 2 fish

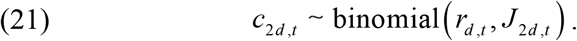

However, protocols for age assignment of older fish varied across years. In years prior to 2017, age 1 or age 2 was assigned based on size thresholds for subsampled groups, *g*, whose lengths were measured. The subsampled groups of the collected fish were pooled over 1-to 6-month periods depending on the year. Because age assignments were based on a measured sub-sample, we accounted for this sampling error by first estimating age fractions from the sub-sample and then applying these fractions to the total catch of fish in the sample tank. First, the total number of age 1 and 2 fish subsampled for a group, *s*_*g*,*t*_, and the number of age 1 fish, *n*_1*g*,*t*_, were used to estimate the fraction of age 1 fish (*p*_1*g*,*t*_):

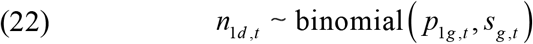

Given that day *d, t*, fell within the period encompassed by group *g, t*, the daily abundance of age 1 juvenile Coho salmon was calculated:

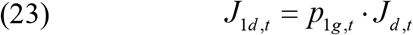

and by subtraction the abundance of age 2 juvenile Coho salmon was calculated:

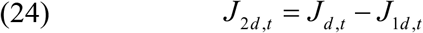

Thus, we assume a constant age fraction over each subsample period. The total annual abundance for each age at outmigration, *o*, was then obtained by adding the *J*_*o*,*d*,*t*_ over the total number of sample and fish collection days in calendar year *t, D*_*t*_:

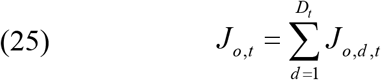

## Results

### Summary of Data Used in the Life Cycle Model

The total escapement of Coho salmon upstream of Swift Dam was variable but increased slightly over time with the lowest escapement recorded in 2015 (3,754 fish) and the highest in 2020 (9,486 fish; fig. 5). The annual escapement of hatchery-origin fish averaged 5,417 (SD=1,486) with the highest number and fraction of hatchery fish escaping in 2014, whereas natural-origin escapement increased steadily over the time series (fig. 6). In 2013 just 28 naturally produced adult Coho salmon were transported upstream of Swift Dam and by 2020 4,863 natural-origin adults were transported.

**Figure 5.**
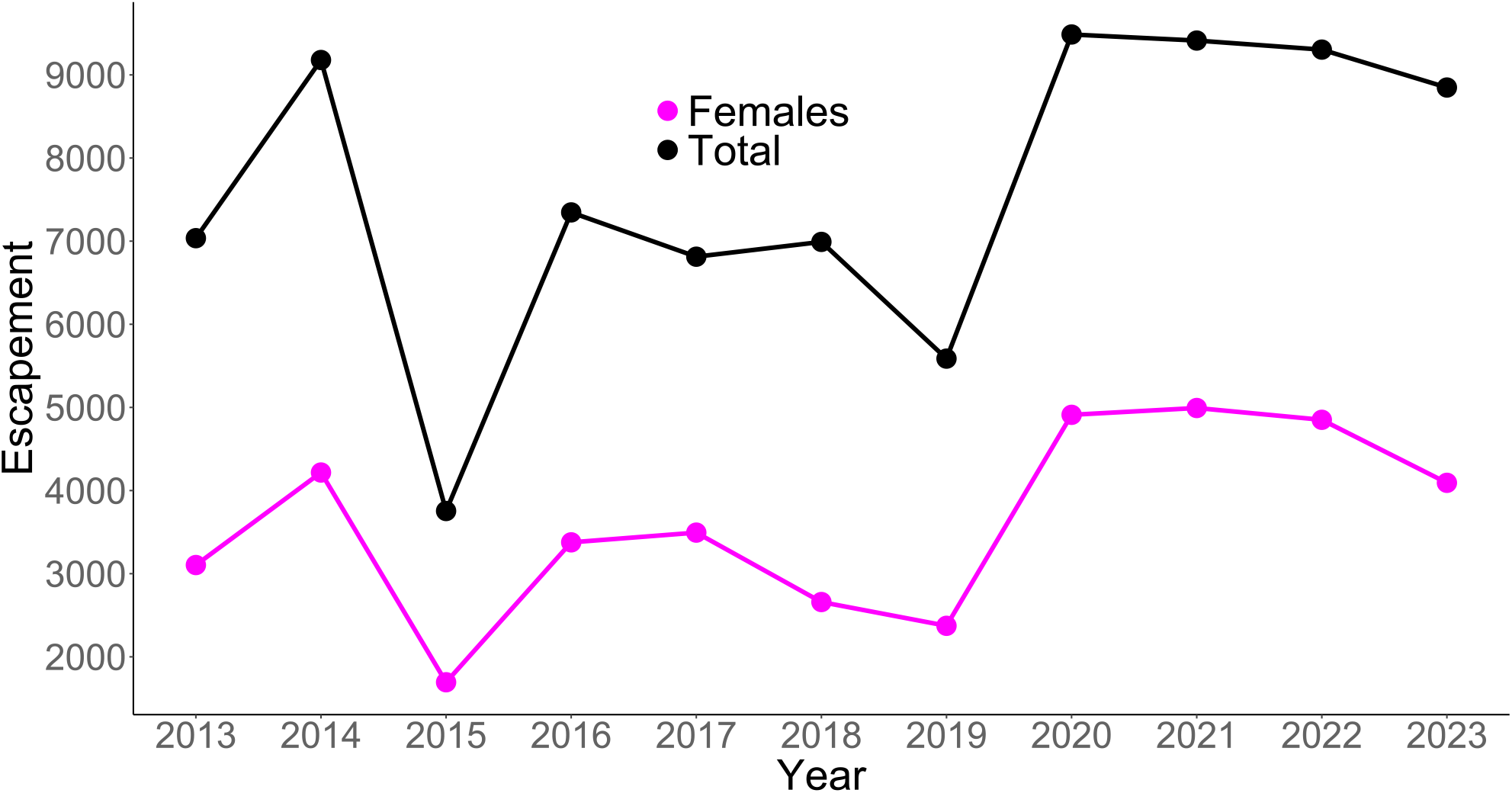
Time series of total and female escapement for Lewis River basin Coho salmon upstream of Swift Dam.

**Figure 6.**
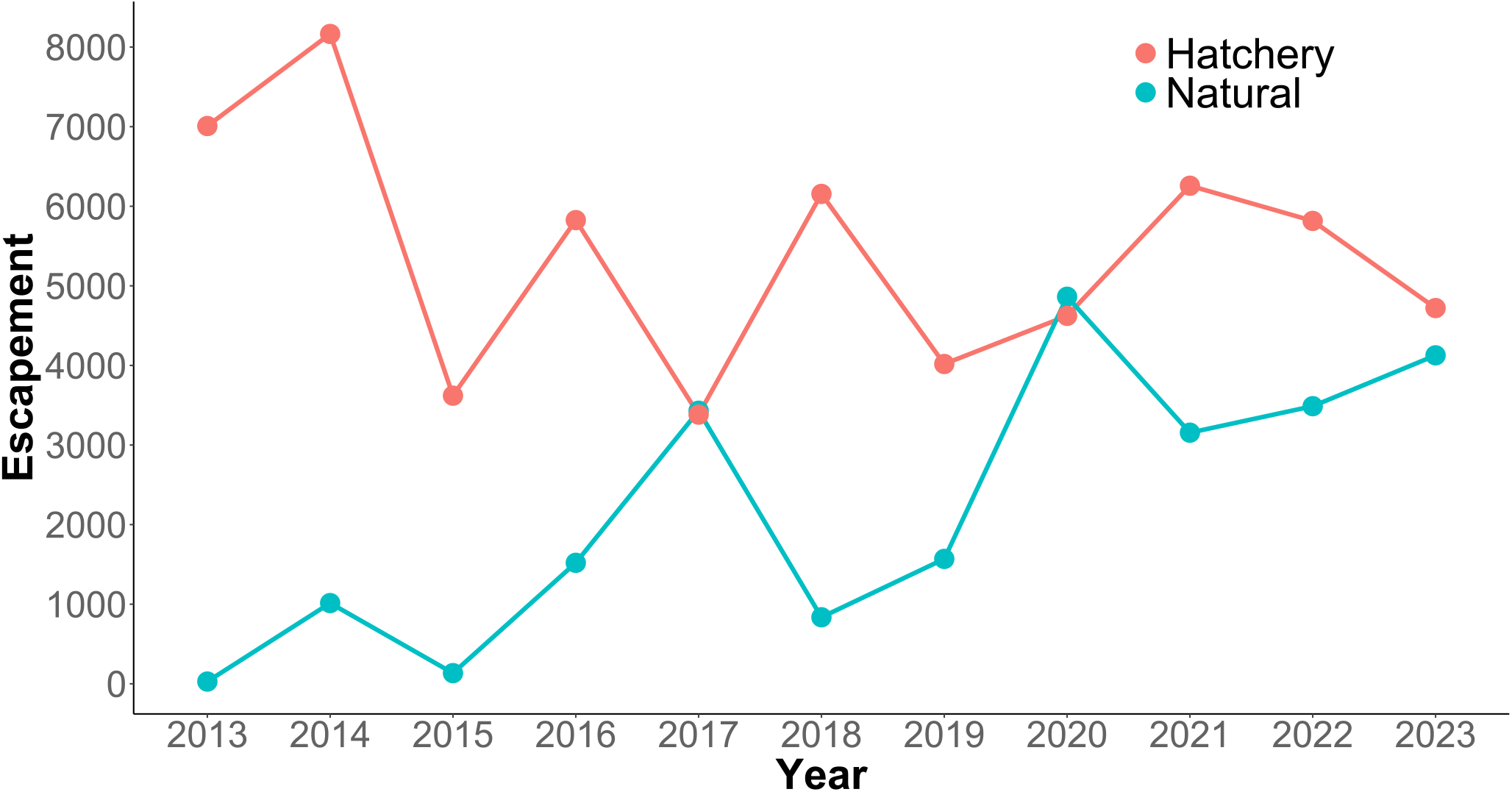
Time series of hatchery and natural -origin escapement for Lewis River basin Coho salmon upstream of Swift Dam.

The proportion of hatchery-origin female spawners (pHOS) generally decreased over time, but pHOS was highly variable, averaging 0.75 (SD=0.182) over the time series. Values of pHOS ranged from a high of 0.99 in 2013 to low of 0.48 in 2017 and 2020 (fig. 7). The proportion of early spawning Coho salmon transported upriver of Swift Dam (*Ep*) was 1.0 during 2013 and 2014, sharply dropping to 0.08 in 2015, and then showing an increasing trend equaling 0.8 in 2023 (fig. 8).

**Figure 7.**
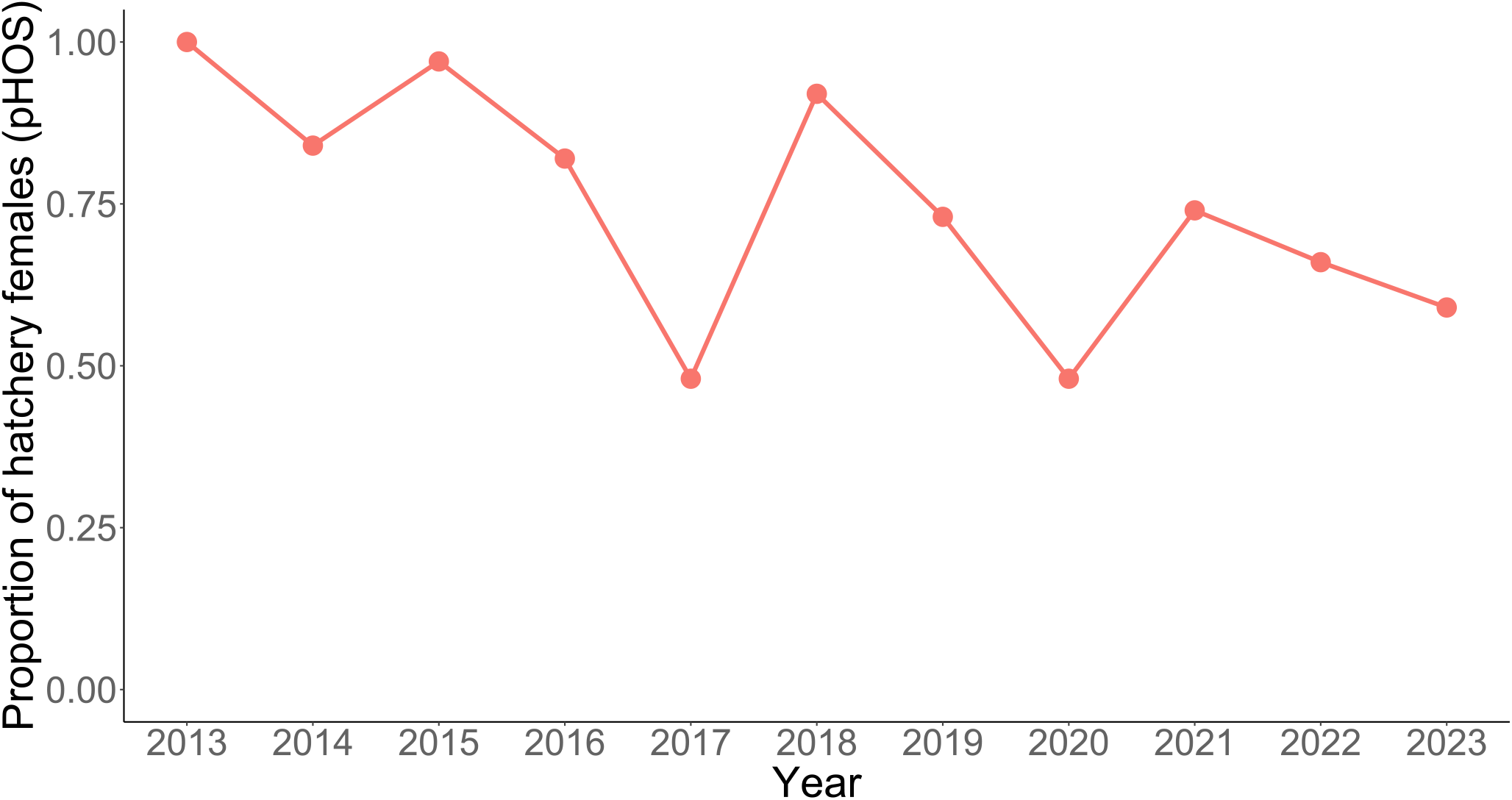
Time series of the proportion of hatchery-origin females out of the total female Coho salmon spawning upstream of Swift Dam in the Lewis River basin.

**Figure 8.**
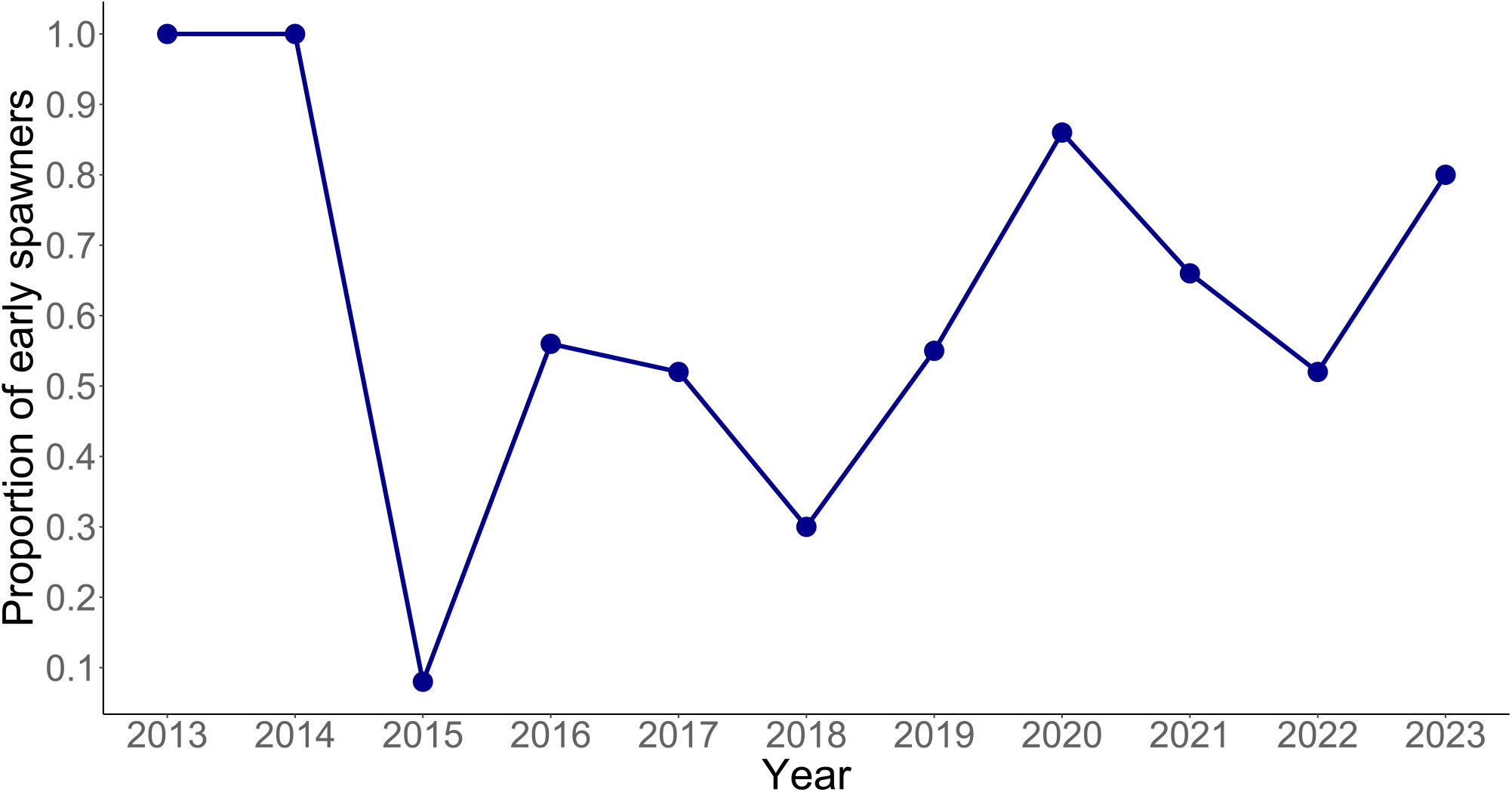
Time series of the fraction of early spawning females out of the total female Coho salmon spawning upstream of Swift Dam in the Lewis River basin.

The number of natural-origin juvenile Coho salmon collected at the Swift FSC (from hatchery and natural-origin spawners) showed an increasing trend over time (fig. 9). Over the annual time series, the average number of juvenile Coho salmon collected at the Swift FSC was 50,896 fish (SD±30,771) and ranged from 9,201 (SD±46) fish in 2014 to 100,137 (SD±766) in 2019.

**Figure 9.**
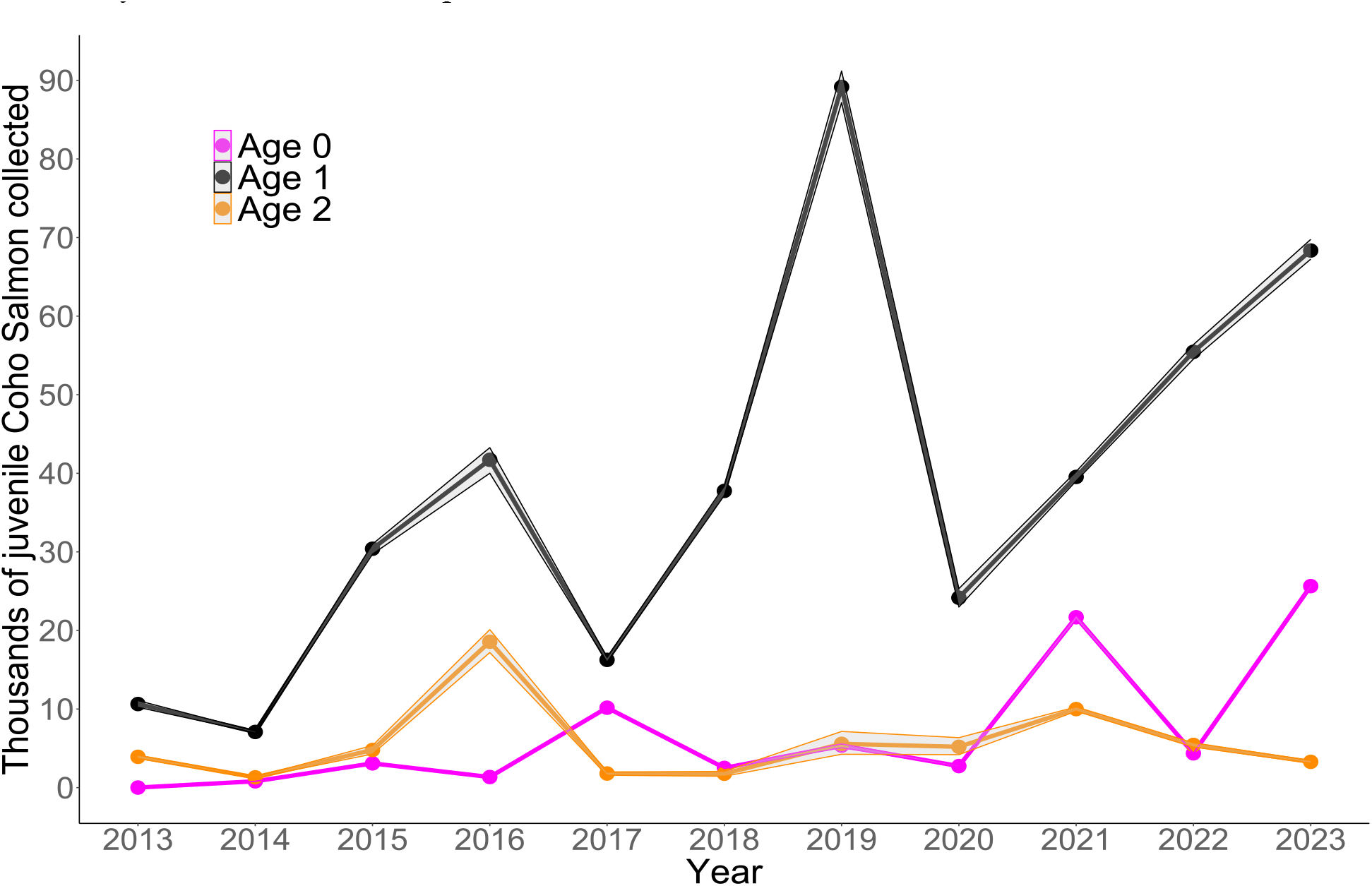
Time series of Lewis River juvenile Coho salmon by age at outmigration collected annually at the Swift FSC. Shaded area represents the 5^th^ and 95^Th^ credible limits.

Calculation of the proportion of juveniles transported by age at outmigration showed over the entire time series, age 1 juveniles always represented a majority (> 0.55) of the fish collected at the Swift FSC each year (fig. 10). However, in some years either age 0 or age 2 fish represented a sizable proportion of the total number of fish transported. For instance, in 2016, age 2 fish represented 30% of all fish collected. Similarly, age 0 fish represented 36% of all collected juveniles in 2017 and 30% in 2021. Thus, even though age 1 fish were predominant in each year, in some years either very young or older fish represent a sizeable portion of the juveniles collected.

**Figure 10.**
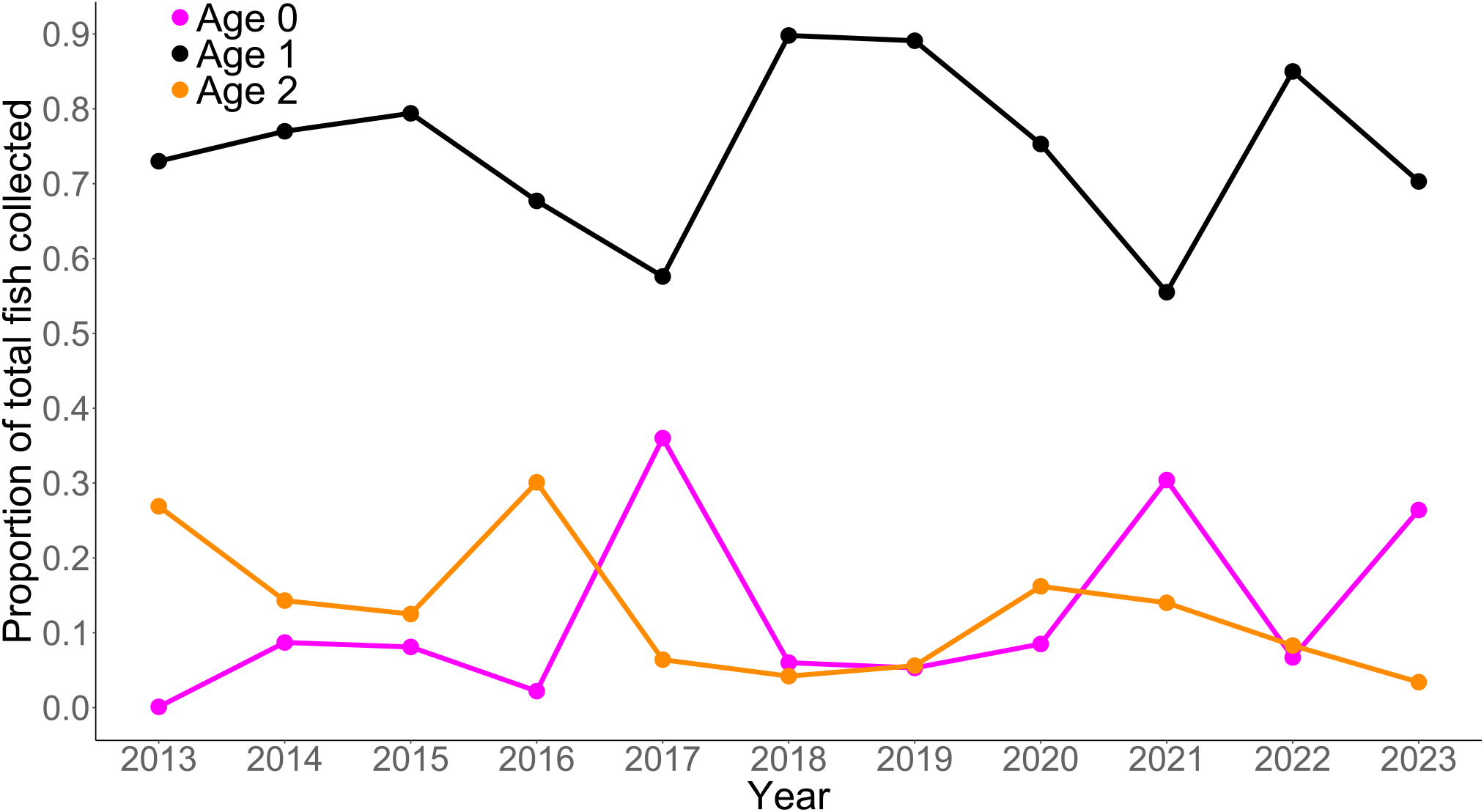
Time series of the fraction of age 0, age 1, and age 2 juvenile Coho salmon collected at the Swift FSC.

The number of juveniles collected at the Swift FSC is determined by the number of spawning females upstream of Swift Dam and the FCE of juveniles at the Swift FSC. Juvenile Coho salmon FCE at Swift Dam averaged 0.375 (SD=0.196) and ranged from a low of 0.06 in 2013 to a high of 0.64 in 2019. The time series of FCE suggests an increasing trend in FCE over time (fig. 11). More recent years (2019 to 2023) had higher FCE (mean=0.536, SD=0.129) than years before 2019 (mean=0.242, SD=0.127; fig. 11). These changes in FCE correspond to improvements made to the Swift FSC since it was commissioned in late 2012 (PacifiCorp, 2024). Because the number of spawning adults and juveniles has increased, both the number of female spawners and the effect of FCE must be accounted for when estimating life cycle productivity.

**Figure 11.**
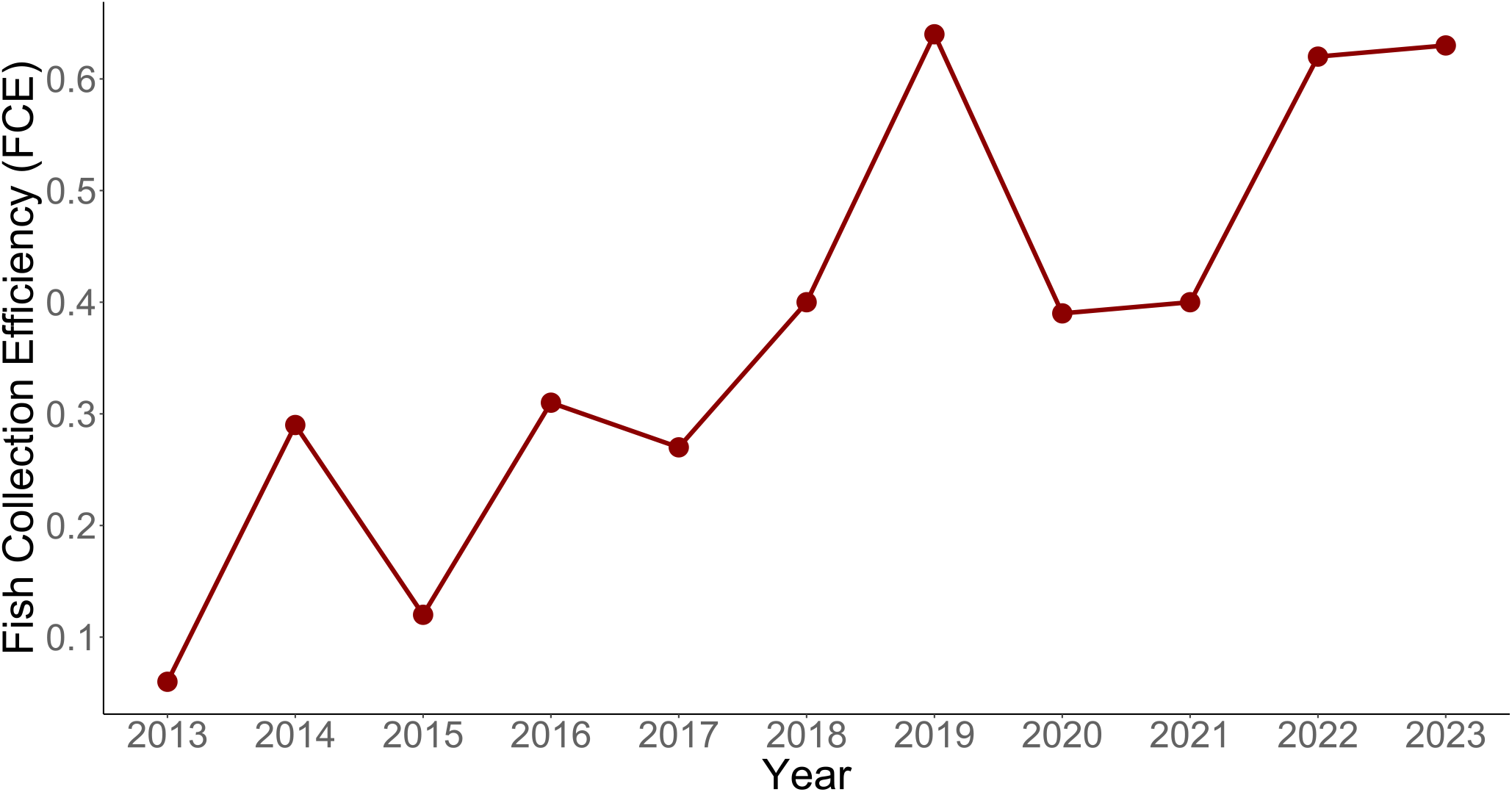
Time series of average annual Fish Collection Efficiency (FCE) at the Swift FSC.

One factor that could also affect the productivity and survival of juvenile Coho salmon upstream of Swift Dam is the minimum summer flows during outmigration. Lower summer flows could result in warmer temperatures and physiological stress, higher predator activity, reduced habitat area and thus, lower fish production at Swift Dam. On average, minimum mean monthly summer flow for our index site on the Muddy River was 149 ft^3^/s (SD=29.4) and ranged from 111 ft^3^/s in 2015 to 209 ft^3^/s in 2017 (fig. 12). In preliminary analyses, we evaluated the effect of maximum winter flows on productivity; however, we found no effect, and to simplify this already complicated first report of the life cycle model, we do not present our results of maximum winter flows.

**Figure 12.**
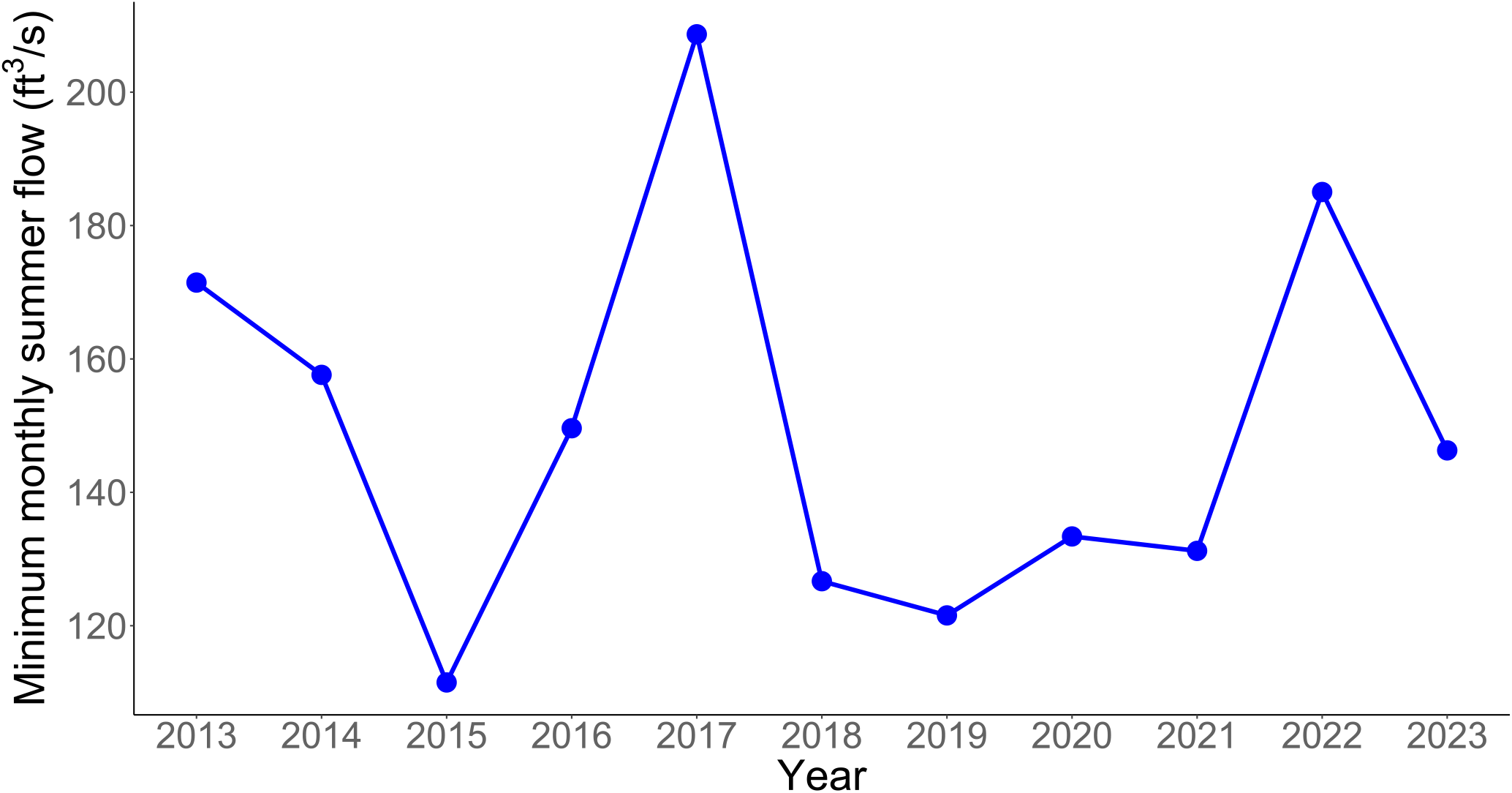
Time series of minimum mean monthly summer (June, July, and August) flow as measured at the Muddy Creek tributary upriver of Swift Reservoir (USGS station number 14216500; U.S. Geological Survey, 2024).

### Parameter Estimates Under the State-Space Model

Posterior distributions represent the uncertainty about the model parameters (table 1; fig. 13 and 14). Annual FCE was found to have a strong positive effect on productivity with a median = 1.962 (95% credible interval (CI) = 0.247 – 4.079). The fraction of the posterior distribution below zero was 0.008, indicating that FCE at the Swift FSC had a positive effect on the number of recruits per spawner that contribute to the population. The median of the posterior distribution for mean productivity across years when FCE = 1 was α_*FCE*=1_= 64 naturally produced juvenile recruits per female spawner (95% CI = 10.37 – 245.79; table 2, fig. 14). Assuming a mean fecundity of 2,600 eggs (Beacham, 1982) per female, this equates to a mean egg-to-juvenile outmigrant survival of about 2.4% at low spawner density.

**Table 1.** Summary statistics for posterior distributions for the two-stage life cycle model of Lewis River Coho salmon showing parameters for the Beverton-Holt model, smolt-to-adult return rate, observation, and process error, as well as the autorcorrelation coefficients. Also shown is the estimated mean productivity should FCE = 1 and capacity when estimated with an informed and weakly informed prior distribution. Notice all estimates are provided while the effect of FCE on juvenile productivity is included in the model.

| Parameter | Lower<br>95% CI | Median | Upper<br>95% CI | Mean | SD |
| --- | --- | --- | --- | --- | --- |
| **Beverton-Holt stock-recruitment model** |  |  |  |  |  |
| $\theta_0$ (intercept) | 1.465 | 2.196 | 2.829 | 2.195 | 0.357 |
| $\theta_1$ (FCE effect) | 0.247 | 1.962 | 4.079 | 2.062 | 0.958 |
| * $\theta_2$ (pHOS effect) | -3.445 | -1.013 | 1.473 | -0.972 | 1.243 |
| * $\theta_3$ (Early effect) | -0.660 | 0.444 | 1.791 | 0.531 | 0.683 |
| * $\theta_4$ (Max winter flow effect) | -0.233 | 0.072 | 0.321 | 0.053 | 0.181 |
| * $\theta_5$ (Min summer flow effect) | -0.590 | -0.332 | 0.071 | -0.301 | 0.185 |
| Productivity at FCE = 1 ($\alpha_{FCE=1}$) | 10.370 | 64.023 | 245.794 | 94.870 | 159.163 |
| $K$ (capacity) informed prior distribution | 84,584 | 297,901 | 812,488 | 361,546 | 237,765 |
| $K$ (capacity) weakly informed prior distribution | 58,373 | 645,779 | 1,903,420 | 781,146 | 564,100 |
| Smolt-to-Adult return rate (SAR) | 0.001 | 0.0449 | 0.130 | 0.058 | 0.064 |
| **Process error** |  |  |  |  |  |
| $\sigma_{P,A}$ (adults) | 1.094 | 1.882 | 3.038 | 1.983 | 0.554 |
| $\sigma_{P,J}$ (juveniles) | 0.084 | 0.217 | 0.410 | 0.231 | 0.093 |
| **Autocorrelation** |  |  |  |  |  |
| $\phi_A$ (adults) | -0.306 | 0.266 | 0.853 | 0.255 | 0.309 |
| $\phi_J$ (juveniles) | -0.999 | -0.511 | 0.431 | -0.415 | 0.466 |

Included as a stand-alone effect in addition to the effect of FCE on productivity.

**Figure 13.**
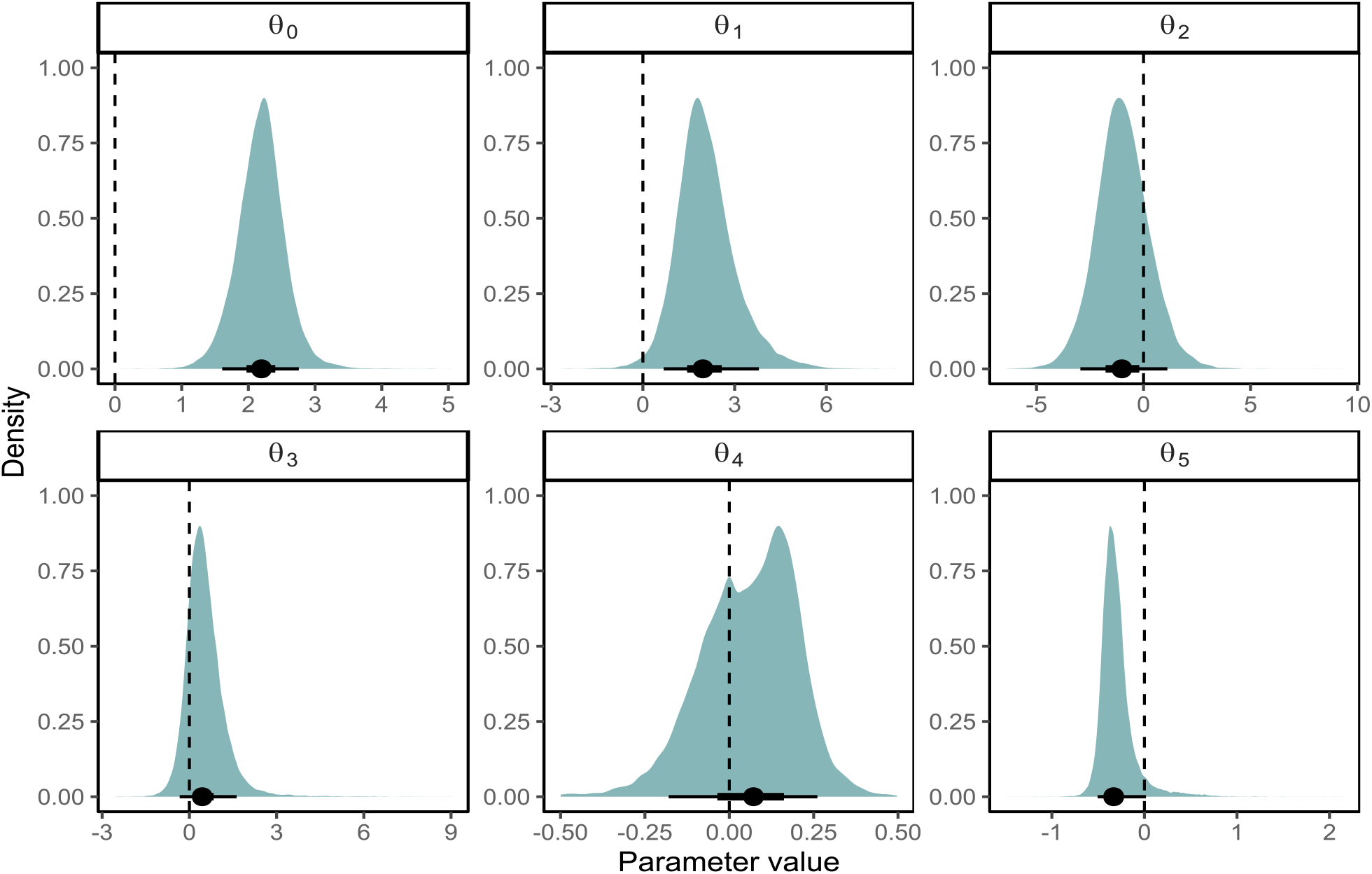
Posterior distributions for the intercept *θ* _0_ and effect of fish collection efficiency (*θ*_1_) on productivity, the effect of the fraction of hatchery female spawners (*θ* _2_) and early spawners (*θ*_3_), and effects of the maximum monthly winter flow (*θ* _4_) and minimum monthly summer flow (*θ*_5_) under the Beverton-Holt stock-recruitment model for Lewis River Coho salmon. Posterior summaries at the bottom of each panel show the median (circle), 90% credible interval (thin line), and 50% credible interval (thick line). Vertical dashed line denotes 0.

**Figure 14.**
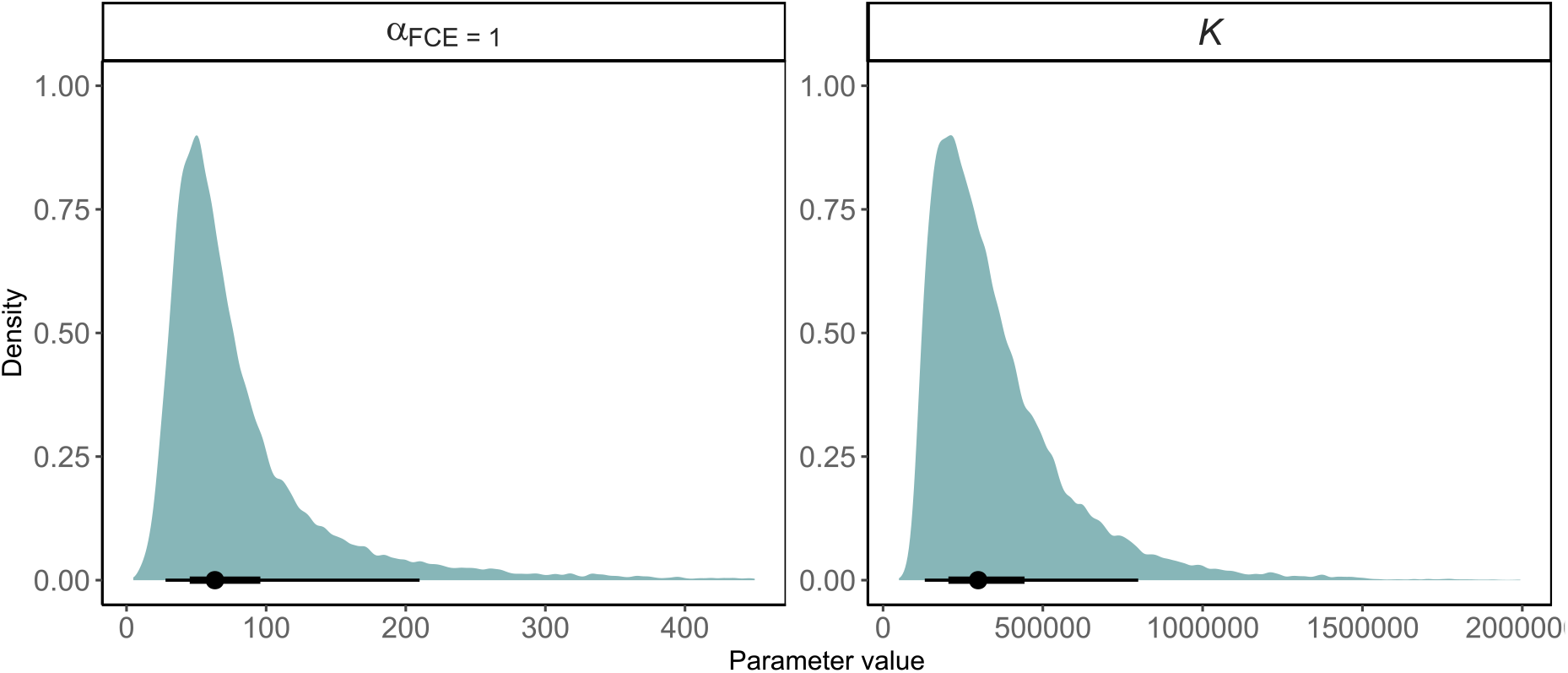
Posterior distributions for productivity when fish collection efficiency (FCE) = 1, and carrying capacity (*K*) as estimated under the Beverton-Holt stock-recruitment model for Lewis River Coho salmon. A prior distribution with parameters from Korman and Tompkins (2014) were used to inform the posterior distributions. Posterior summaries at the bottom of each panel show the median (circle), 90% credible interval (thin line), and 50% credible interval (thick line)

Evaluations of individual effects on productivity over and above the effects of FCE indicated that minimum monthly summer flow had a negative effect on annual productivity (table 1; fig. 13). About 95% of the posterior distribution for *θ*_5_ were < 0, indicating statistical support for an effect of minimum summer flow on productivity. Also, the fraction of hatchery spawners tended to have a negative effect on productivity with 80% of the posterior distribution having *θ* _2_ < 0. In contrast, a larger proportion of early spawning Coho salmon was associated with generally higher juvenile productivity with 17.5% of the posterior distribution having *θ*_3_ < 0. The effect of maximum monthly winter flows on juvenile productivity was not statistically supported with a posterior distribution that encompassed zero with 34% of the distribution below zero.

Using population level estimates of capacity from Korman and Tompkins (2014) to inform the prior distribution for capacity (*K*) of Lewis River Coho salmon resulted in a well-defined posterior distribution with a median capacity of 297,901 (95% CI = 84,584 – 812,488) maximum juveniles produced (table 1; fig. 14 and 15). Uncertainty about the estimate of capacity is demonstrated by how the choice in prior distribution can influence the estimate of capacity. For example, for a weakly informed prior distribution we used a truncated normal distribution where *K* ∼ *normal*(*μ* =10,000, SD =1,000,000), median capacity was 645,779 (95% CI = 58,373 – 1,903,420) juveniles, and the posterior distribution was much more protracted compared to the posterior distribution informed by Korman and Tompkins (2014; fig. 14), *K* ∼ *lognormal*(*μ* =7.34, SD=0.67). Nonetheless, the posterior distributions were similar to their respective prior distributions, supporting the conclusion that the data provided little information towards the estimation of capacity in the life cycle model.

**Figure 15.**
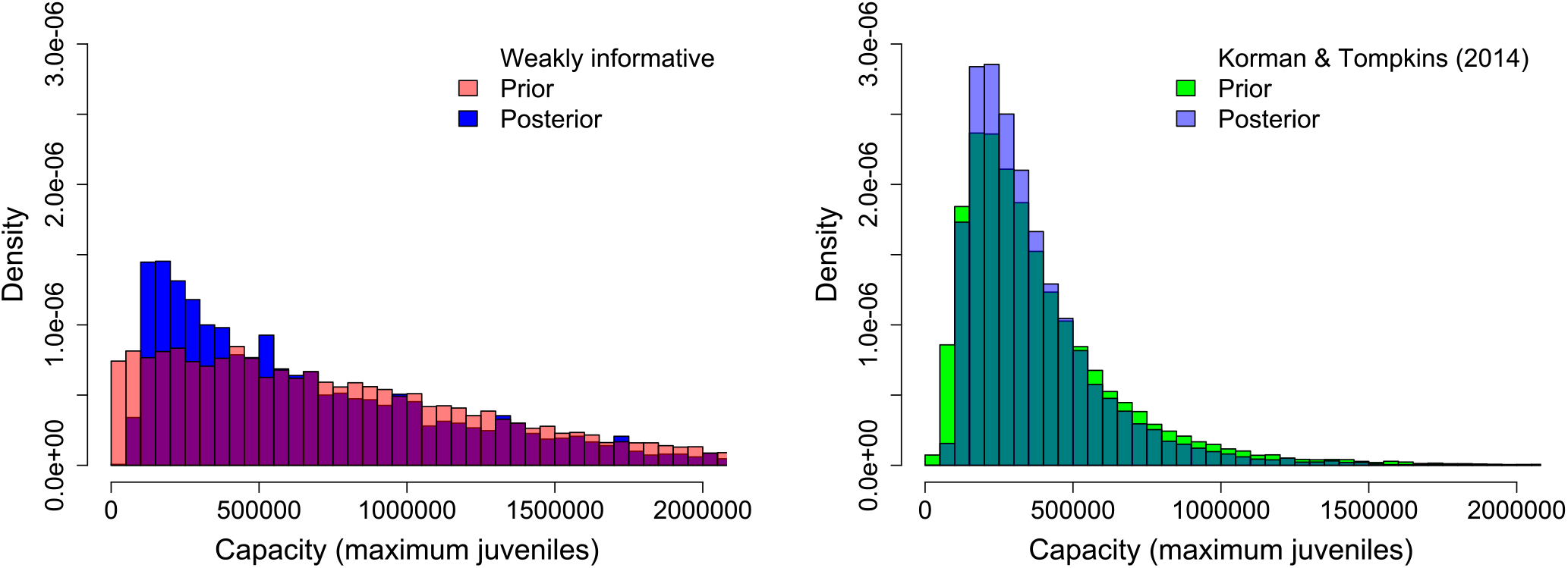
Comparison of using a weakly informed (left panel) versus an informed (right panel) prior distribution in relation to their respective posterior distributions for estimating carrying capacity of Lewis River Coho salmon upriver of Swift Dam.

The curves from the Beverton-Holt stock-recruitment model illustrate how recruitment of naturally produced juveniles varies as a function of the number of female spawners (fig. 16). The data used to inform the stock-recruitment model encompassed a relatively narrow range in the number of spawners and juveniles – from a low of 1,074 to about 4,993 female spawners. Therefore, our estimated stock-recruitment model parameters provide inference over a narrow range of observed population levels. This is also evidenced by the uncertainty in our estimates of capacity (fig. 15).

**Figure 16.**
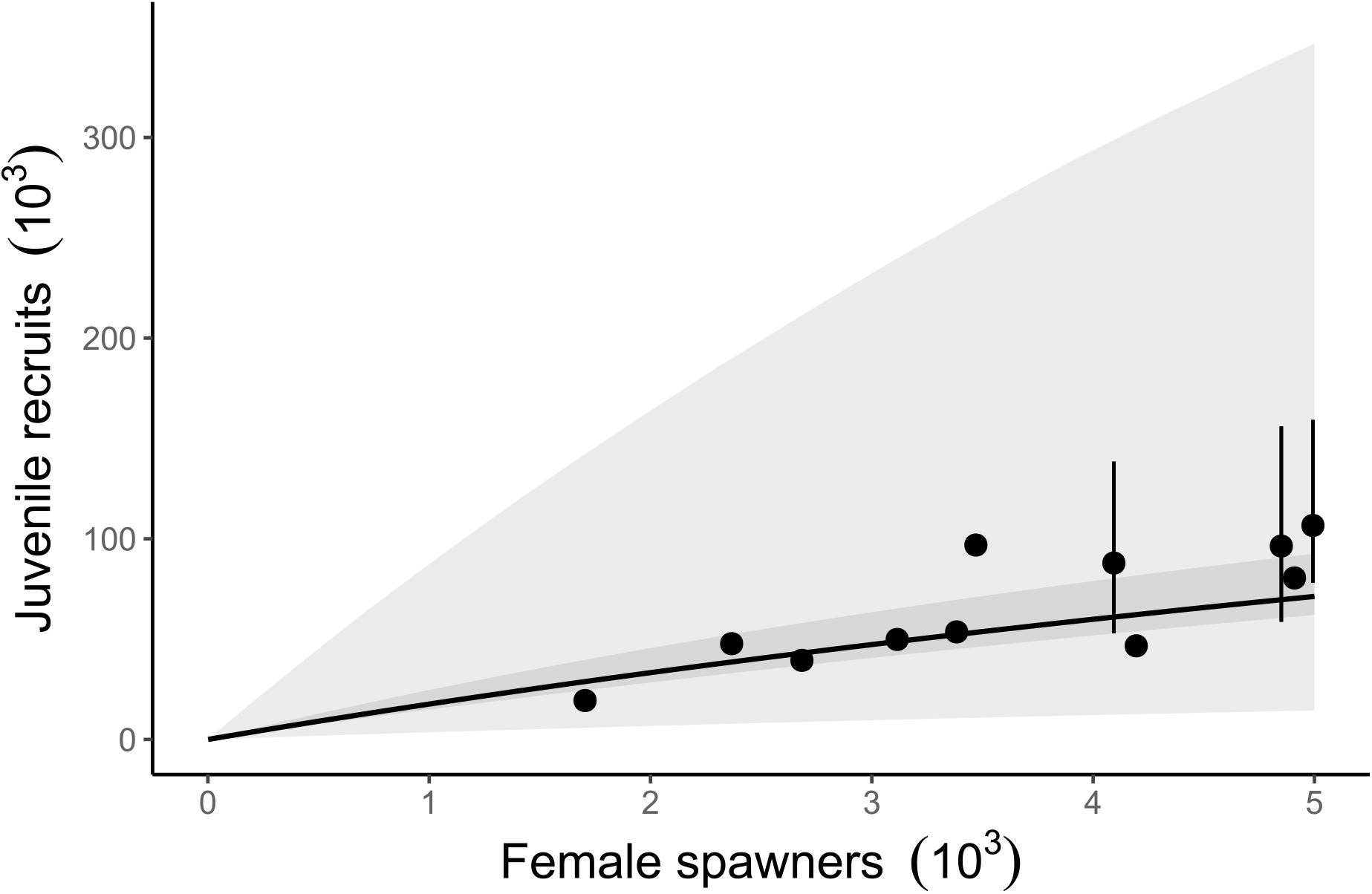
The fitted Beverton-Holt function based on posterior distribution medians of parameters expressed as production of juvenile recruits. Note the stock recruitment curve was estimated using the mean FCE = 0.375 over the time series. Dark shaded area represents the 95% credible about the mean trend and light shaded area represents the 95% credible interval when process error is considered.

For the juvenile-to-adult transition, the median of the posterior distribution for the mean SAR across all years was 0.045 (95% CI = 0.001 – 0.13; table 1). Thus, in the absence of accounting for harvest, a median of 4.5% of juvenile outmigrants returned to the Lewis River. Owing to the hierarchical structure of the state-space model, annual variation in productivity and SAR can be estimated as random effects drawn from lognormal process error (*σ* _P,J_ and *σ* _P,A_, see Table 2; fig. 16, 17). Both SAR and productivity vary considerably over time and generally increase after brood year 2015. The higher productivity since brood year 2015 coincides with increasing FCE since 2015 (fig. 11). The number of juveniles collected at Swift Dam given the number of female spawners showed the increase in productivity of natural juveniles upstream of Swift Dam because of higher juvenile fish collection performance at the Swift FSC (fig. 16 and 18).

**Figure 17.**
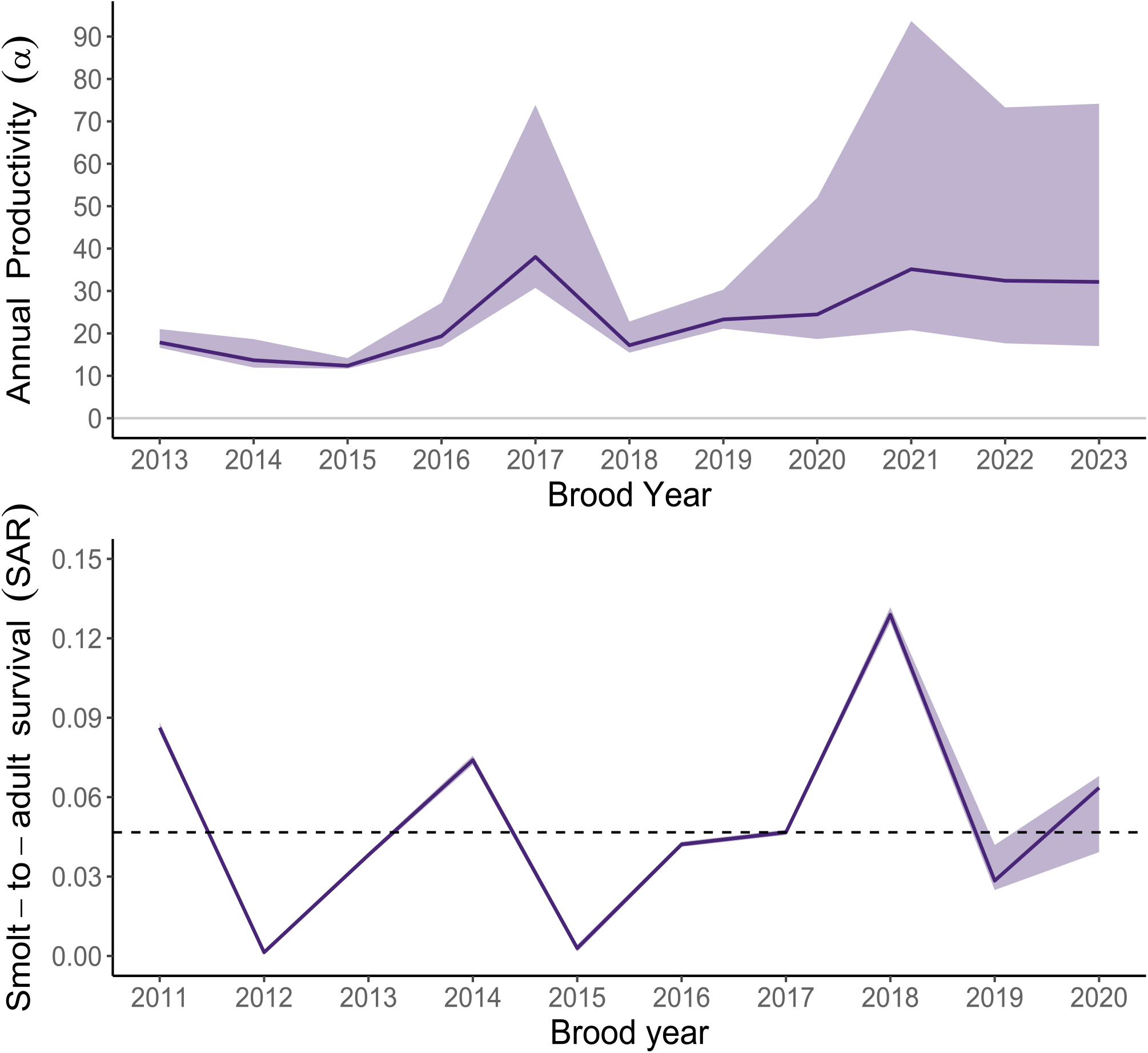
Annual estimates of juvenile productivity (recruits per spawner; top panel) and smolt-to-adult return rate (SAR, bottom panel). Lines connect annual medians of posterior distributions, shaded area represent the 5^th^- 95^th^ credible interval, and the dashed line shows the overall median SAR = 0.045.

**Figure 18.**
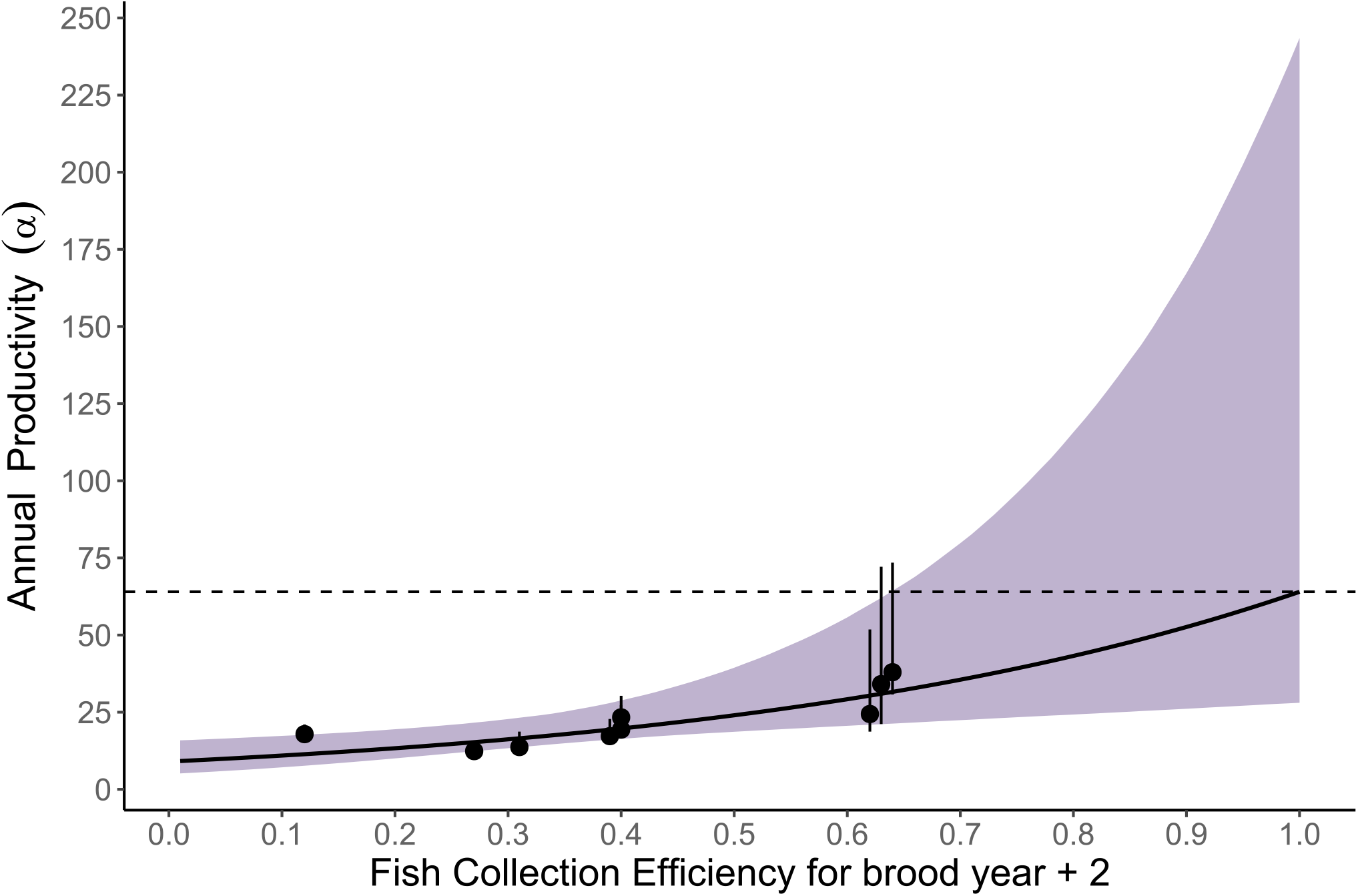
Relationship between fish collection efficiency (FCE) and the number of juveniles per spawner, shaded area represents the 95% credible interval about the mean effect of FCE and the error bars represent the 95% credible interval about the estimated annual productivity. The dashed line represents the extrapolated estimate of productivity when FCE = 1 (pre-collection productivity).

## Discussion

The state-space life cycle model for Lewis River Coho salmon is a model of intermediate complexity but can be used to gain insight about population dynamics and factors affecting different life stages as part of the ongoing reintroduction program. The state-space formulation of the model accounts for both observation and process error, provides estimates of uncertainty in both abundance and demographic parameters, and quantifies contributions of age and sex to the aggregate population. Our formulation of the state-space model provides a wealth of information about population dynamics such as estimates of process uncertainty, carrying capacity and density dependence, juvenile-to-adult survival rates, and factors affecting production of juvenile Coho salmon upstream of Swift Dam, the latter being particularly important for understanding how fish collection efforts at the Swift FSC affect population sustainability and recovery. We also demonstrate how the model can be expanded to include juvenile age at outmigration and subsequent age at return, harvest, as well as ocean effects on demographic parameters such as juvenile to adult return rate.

As described above, the life cycle model for Lewis River Coho salmon can be used to provide estimates of the effect of FCE on adult-to-juvenile productivity. This analysis represents an initial effort that illustrates the utility of multistage statistical life cycle models for understanding the interplay between population dynamics and trap and haul programs that seek to sustain salmon populations. There are many other aspects of the dynamics of this population that we have yet to explore. For example, there is much uncertainty in the estimation of capacity under the Beverton-Holt model because of the limited range in escapement and the number of spawning females. Regardless of the uncertainty, our informed estimates of capacity (median = 297,901) were much larger than recent juvenile fish collection at the Swift FSC, supporting the conclusion that higher escapement numbers might result in higher juvenile abundance. Therefore, firm conclusions regarding the capacity of habitat for Coho salmon upstream of Swift Dam cannot be made at this time. Additional information regarding productivity at higher escapement is needed before a more definitive conclusion can be made. By also incorporating a fuller set of covariates, harvest rates, and juvenile age structure in relation to their age at adult return into the model, parameter estimates may provide a broader set of inferences to the individual contribution of these factors to the life cycle of Coho salmon upstream of Swift Dam.

We estimated the effect of several covariates on juvenile production. We found strong statistical support for a negative effects of high flows on juvenile Coho salmon productivity, but the mechanisms behind such an effect are unclear. High flows may have pushed fish into the reservoir and caused them to experience a higher predation rate. For other effects on juvenile productivity, higher fractions of hatchery fish were moderately associated with lower juvenile productivity with 80% of the posterior distribution having *θ* _2_ < 0, suggesting lower juvenile production at higher fractions of hatchery escapement. Also, higher fractions of early spawning females were poorly associated with higher juvenile productivity with 82.5% of the posterior distribution had *θ* _3_ > 0. After including FCE in the model, there was little variation to be explained from 10 data points, and high uncertainty in model parameters limits the ability to make inferences about model covariates. Additional years of data could help provide a clearer picture of the factors that affect Coho salmon juvenile production upriver of Swift Dam.

There are several significant updates that could improve model fit and provide greater inference for fishery management. These updates include: (1) accounting for ocean harvest rates for naturally produced fish, (2) obtaining data on age at return for each juvenile age at outmigration for more precise indexing of covariates and the estimation of SAR, and (3) evaluating other covariates to explain variation in adult-to-juvenile productivity (a) and smolt-to-adult return rates (SAR; Logerwell and others, 2003).

Some of the future covariates that could help explain variation in adult-to-juvenile production include:

– Using the median date of adult transportation as an alternative measure of spawn timing effects

Future covariates that may help explain variation in annual smolt-to-adult return rates:

– Juvenile age at outmigration
– Pacific Decadal Oscillation (PDO)
– Coastal Upwelling Index (CUI)
– Sea Surface Temperatures (SST)
– Harvest estimates in the ocean and freshwater downriver of Merwin Dam

Including and evaluating these additional covariates within the two-stage life cycle model could greatly improve the inferences drawn about the production and viability of Lewis River Coho salmon.

Regardless of the improvements that can be made to the model, we can make some reasonable conclusions. First and perhaps foremost, we were able to quantify how FCE at the Swift FSC directly determines the productivity of juveniles. This was demonstrated by the relationship between mean annual productivity and FCE (fig. 18). Thus, the annual progression towards higher FCE at the Swift FSC (i.e., since 2013) contributed to the higher productivity (fig. 17) and greater numbers of juveniles collected in recent years (fig. 9). However, there are few data points at high FCE values (> 0.64) with high uncertainty in juvenile productivity at the upper end of the FCE curve, such as when FCE = 1 (table 1).

Perhaps the model’s most uncertain parameter for the adult-to-juvenile life stage is carrying capacity of the habitat upriver of Swift Dam, *K*. Over the historical time series, there have not been enough spawners released upstream of Swift Dam to elicit compensation in juvenile fish production. Consequently, there is much uncertainty in our estimation of Coho salmon capacity for the habitat upstream of Swift Dam. This is evident when comparing the results from using an informed and uninformed prior distribution to estimate capacity. Using the distribution parameters from the meta-analysis of Coho salmon capacity provided by Korman and Tompkins (2014) as an informed prior distribution yielded a much lower and certain estimate of capacity than using a weakly informed prior distribution, which yielded a much higher and uncertain estimate of capacity (table 1). Both posterior distributions closely followed their respective prior distribution, which suggests the data alone are insufficient for estimating capacity. In preliminary model runs, we compared median capacity with and without the 14.5 km distance of Swift Reservoir, and the difference was trivial when considering the uncertainty in the capacity estimate, with the distance of the reservoir producing an additional 21,447 juveniles. However, this assumes the reservoir and riverine habitat upriver of the reservoir provide the same production and capacity benefits for Coho salmon. Future efforts towards releasing a sufficiently high number of adult Coho salmon upstream of Swift Dam will help inform our estimate of capacity and when compensation in juvenile production would be expected.

The median SAR over all estimable years of about 4.5% is relatively high when considering that this survival rate does not account for harvest mortality. Estimates of SAR by brood year ranged from 0.1% in 2012 to 12.9% in 2018 (fig. 17). However, we should be cautious in drawing much inference from our current estimates of SAR because the model does not yet account for juvenile outmigration and their respective age at return, which can bias age-averaged estimates of SAR. Analysis of scale samples from returning adults with both juvenile age at ocean entry and adult age at return could be incorporated into the model to help resolve this uncertainty. Notwithstanding this limitation, our annual estimate of SAR suggests that in some years, return rates may be high enough to meet replacement and sustain the population so long as juvenile production and fish collection remain at some yet unknown but sufficiently high value. Future applications of the model could include a population viability analysis to determine what FCE target may be needed to sustain Lewis River Coho salmon.

Ultimately, fisheries managers need tools to help understand how potential management actions affect population abundance at different parts of the life cycle to assess whether recovery goals are being achieved. The state-space life cycle model for Lewis River Coho salmon takes a first step towards development of such a tool. Additional work could help better account for key management factors affecting the population (e.g., harvest) to quantify the contribution of the trap-and-haul program performance and incorporate effects of environmental variation. Because the model incorporates unaccounted for variability in demographic parameters, we can use the model to simulate stochastic population trajectories that include important management “knobs” (e.g., FCE, harvest) under expected environmental variation. Such a model would allow managers to assess the relative effect of alternative management scenarios on population trajectories in a probabilistic manner to understand the likelihood of success of a given management action in the face of environmental variability and uncertainty.

## Appendix

**Table A1.** Summary statistics for posterior distributions of adult-to-juvenile productivity for Lewis River Coho salmon estimated for each brood year.

| Parameter | Lower 95% CI | Median | Upper 95% CI | Mean | SD |
| --- | --- | --- | --- | --- | --- |
| $\alpha$ [2013] | 16.25 | 17.86 | 21.11 | 18.28 | 10.258 |
| $\alpha$ [2014] | 11.43 | 13.68 | 18.75 | 14.37 | 25.229 |
| $\alpha$ [2015] | 11.48 | 12.36 | 14.24 | 12.61 | 55.983 |
| $\alpha$ [2016] | 16.24 | 19.34 | 27.34 | 20.47 | 28.661 |
| $\alpha$ [2017] | 28.62 | 38.02 | 74.19 | 43.85 | 11.660 |
| $\alpha$ [2018] | 14.93 | 17.20 | 22.84 | 18.01 | 13.445 |
| $\alpha$ [2019] | 20.45 | 23.29 | 30.46 | 24.30 | 5.175 |
| $\alpha$ [2020] | 17.06 | 24.47 | 52.18 | 28.89 | 4.952 |

**Table A2.** Summary statistics for posterior distributions of Juvenile-to-Adult return rates (SAR) for Lewis River Coho salmon showing parameters for each brood year.

| Parameter | Lower 95%<br>CI | Median | Upper 95% CI | Mean | SD |
| --- | --- | --- | --- | --- | --- |
| SAR overall | 0.0451 | 0.00136 | 0.13 | 0.0578 | 0.063 |
| SAR [2011] | 0.083 | 0.086 | 0.089 | 0.086 | 0.001 |
| SAR [2012] | 0.001 | 0.001 | 0.002 | 0.001 | 0.000 |
| SAR [2013] | 0.037 | 0.038 | 0.040 | 0.038 | 0.001 |
| SAR [2014] | 0.071 | 0.074 | 0.077 | 0.074 | 0.001 |
| SAR [2015] | 0.001 | 0.003 | 0.005 | 0.003 | 0.001 |
| SAR [2016] | 0.041 | 0.042 | 0.044 | 0.042 | 0.001 |
| SAR [2017] | 0.045 | 0.047 | 0.048 | 0.047 | 0.001 |
| SAR [2018] | 0.124 | 0.129 | 0.133 | 0.129 | 0.002 |
| SAR [2019] | 0.022 | 0.028 | 0.049 | 0.031 | 0.008 |
| SAR [2020] | 0.026 | 0.064 | 0.070 | 0.058 | 0.013 |
| SAR [2021] | 0.000 | 0.060 | 0.498 | 0.126 | 0.158 |

## Abbreviations

FCE: Fish Collection Efficiency
FSC: Floating Surface Collector
SAR: smolt-to-adult return rate
USGS: U.S. Geological Survey
MCMC: Markov Chain Monte Carlo

